# A GRAB sensor reveals activity-dependent non-vesicular somatodendritic adenosine release

**DOI:** 10.1101/2020.05.04.075564

**Authors:** Zhaofa Wu, Yuting Cui, Huan Wang, Kun Song, Zhengwei Yuan, Ao Dong, Hao Wu, Yi Wan, Sunlei Pan, Wanling Peng, Miao Jing, Min Xu, Minmin Luo, Yulong Li

## Abstract

The purinergic signaling molecule adenosine (Ado) modulates many physiological and pathological brain functions,but its spatiotemporal release dynamics in the brain remains largely unknown. We developed a genetically encoded <u>G</u>PC<u>R</u>-<u>A</u>ctivation–<u>B</u>ased <u>Ado</u> sensor (GRAB_Ado_) in which Ado-induced changes in the human A_2A_ receptor are reflected by fluorescence increases. This GRAB_Ado_ revealed that neuronal activity-induced extracellular Ado elevation was due to direct Ado release from somatodendritic regions of the neuron, requiring calcium influx through L-type calcium channels, rather than the degradation of extracellular ATP. The Ado release was slow (∼30 s) and depended on equilibrative nucleoside transporters (ENTs) rather than conventional vesicular release mechanisms. Thus, GRAB_Ado_ reveals an activity-dependent slow Ado release from somatodendritic region of the neuron, potentially serving modulating functions as a retrograde signal.

## Main text

Extracellular adenosine (Ado) plays an important role in a wide range of physiological processes (*1-3*), including the sleep-wake cycle, learning and memory, cardiovascular function, and immune responses. Moreover, impaired adenosinergic signaling has been implicated in a variety of diseases and conditions (*2, 4*) such as pain, migraine, epilepsy, stroke, drug addiction, and neurodegeneration (e.g., Parkinson’s disease). In the brain, Ado acts as a neuromodulator or a homeostatic modulator at the synaptic level, through the activation of distinct G protein-coupled Ado receptors (*5*). Although Ado’s function has been extensively studied, the mechanisms underlying Ado release in the brain remains poorly understood, the difficulty lies largely in the lack of sensitive methods for directly detecting Ado *in vivo* with both cell-type specificity and high spatiotemporal resolution. Recently, a group of genetically encoded <u>G</u>PC<u>R</u>-<u>A</u>ctivation–<u>B</u>ased (GRAB) sensors were developed for measuring the dynamics of several neuromodulators—including acetylcholine, dopamine, and norepinephrine—and had been used under various *in vivo* conditions (*6-9*). Using a similar strategy, we designed a genetically encoded GRAB sensor for Ado (Fig. 1A) and used this novel tool to examine the mechanism underlying a neuronal activity–dependent Ado release.

**Fig. 1.**
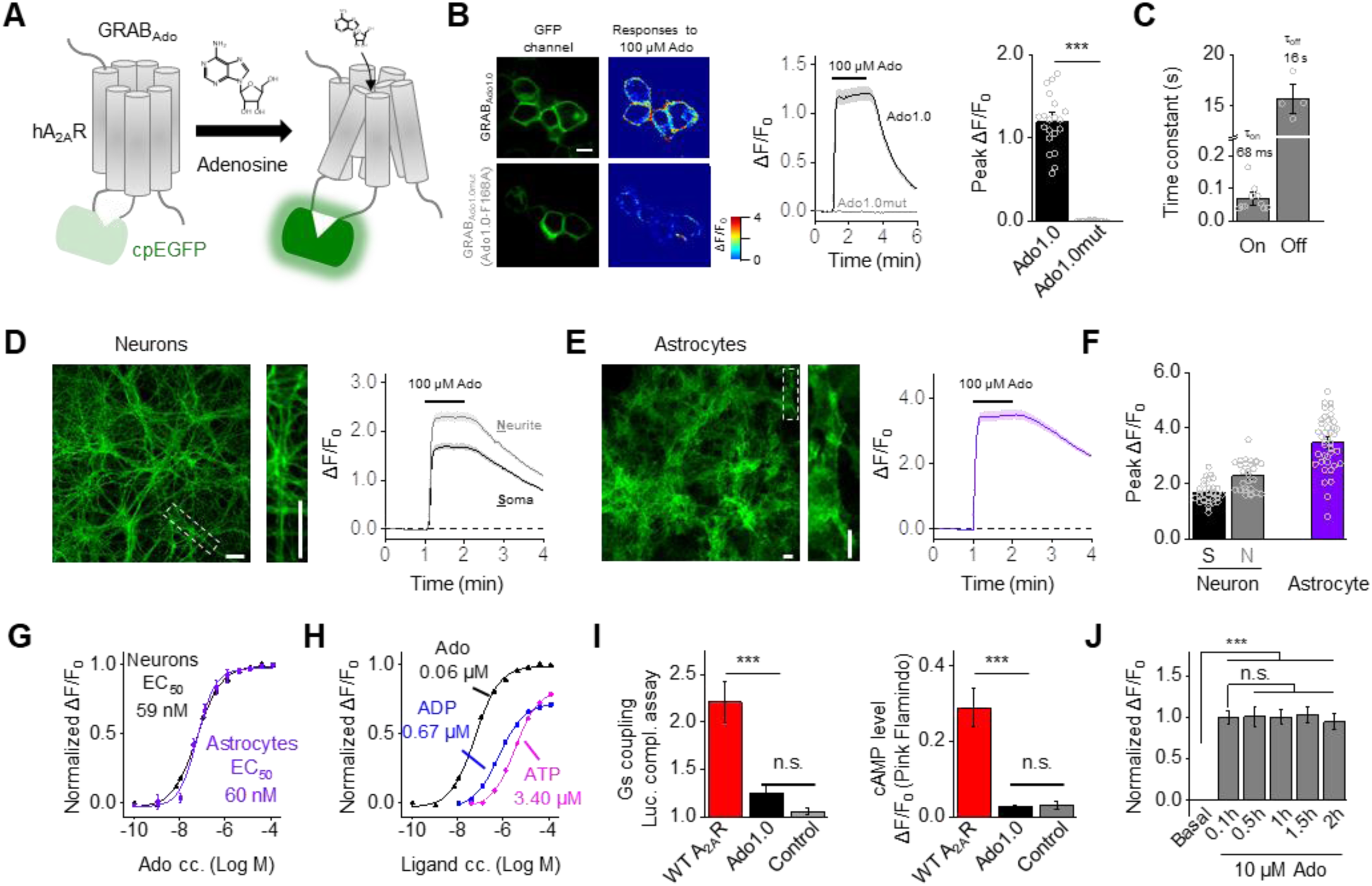
Design and characterization of genetically encoded fluorescence-based adenosine sensors. **(A)** Schematic drawing depicting the principle of the GRAB-based Ado sensors designed using the human A_2A_R as the scaffold combined with circularly permuted enhanced GFP (cpEGFP). Binding of the ligand adenosine induces a conformation change that increases the fluorescence signal. **(B)** Expression, localization, and fluorescence response of the GRAB_Ado1.0_ (Ado1.0) and GRAB_Ado1.0mut_ (Ado1.0mut) sensors in HEK293T cells. Left, representative images of sensor fluorescence before and after application of 100 μM Ado (scale bar, 10 μm). Middle and right, time course and summary of peak ΔF/F_0_ measured in cells expressing Ado1.0 or Ado1.0mut; where indicated, 100 μM Ado was applied; n = 20 cells from 2 cultures. **(C)** Average rise time (τ_on_) and decay time (τ_off_) constants of the change in Ado1.0 fluorescence in response to 100 μM Ado followed by the A_2A_R antagonist SCH-58261 (200 μM); n = 10 and 4 cells, respectively. **(D-F)** Ado1.0 was expressed in cultured neurons **(D)** and astrocytes **(E)**, and Ado was applied where indicated (Scale bars, 30 μm). **(F)** Summary of peak ΔF/F_0_ measured in the soma and neurites (left) and astrocytes (right); n = 28-40 regions of interest (ROIs) measured in 2-3 cultures. **(G)** Normalized dose-response curves for Ado1.0-expressing neurons and astrocytes; n ≥ 30 ROIs from ≥1 culture. **(H)** ΔF/F_0_ was measured in Ado1.0-expressing neurons in response to Ado, ADP, and ATP and normalized to the response measured with 10 μM Ado; n ≥ 20 ROIs each. The data from the Ado group is reproduced from **(G)**. **(I)** Ado1.0 does not engage downstream Gs protein signaling. A luciferase complementation assay was performed to measure Gs protein coupling (left), and the cAMP sensor PinkFlamindo was used to measure cAMP levels (right) in HEK293T or HeLa cells expressing A_2A_R or Ado1.0 (non-transfected cells were used as a control); n ≥ 3 independent experiments each. **(J)** Normalized ΔF/F_0_ was measured in Ado1.0-expressing neurons in response to 10 μM Ado continuously applied over a 2-hour period; n = 28 neurons from 3 cultures.

### Development and characterization of GRAB sensors for adenosine

To develop the genetically encoded GRAB sensor for Ado, we first screened candidate GPCR scaffolds by inserting cpEGFP in the receptor flanked by short linker peptides at both the N- and C-terminus (fig. S1A); we then selected an A_2A_ receptor (A_2A_R)-based chimera (GRAB_Ado0.1_) for further optimization, based on its membrane trafficking and high fluorescence response upon Ado application (fig. S1B). By systematically optimizing the length and amino acid composition of the linkers between the A_2A_R and the cpEGFP, we identified the protein with the largest fluorescence response (fig. S1C), and named it GRAB_Ado1.0_ (hereafter referred to as Ado1.0). When expressed in HEK293T cells, Ado1.0 trafficked to the cell membrane and produced a 120% peak ΔF/F_0_ in response to saturated concentration of extracellular Ado (Fig. 1B); in contrast, a non-ligand-binding mutant form of the sensor (GRAB_Ado1.0mut_, or Ado1.0mut for short) failed to respond to Ado (Fig. 1B and fig. S2E). Finally, Ado1.0 has rapid response kinetics, with a rise time (τ_on_) of 68±13 ms (Fig. 1C)

When expressed in neurons, Ado1.0 was widely distributed throughout the plasma membrane, including the soma, axons, and dendrites (Fig. 1D and fig. S2A). Similarly, when expressed in astrocytes under the control of the GfaABC1D promoter (*10*), Ado1.0 was widely distributed throughout the plasma membrane, including the soma and processes (Fig. 1E). Both neuronal and astrocytic Ado1.0 responded to Ado application in a dose-dependent manner, with similar EC_50_ values of about 60 nM (Fig. 1F and G). In addition, the Ado-induced fluorescence response was blocked by the A_2A_R antagonist SCH-58261 (fig. S2C). We also measured the selectivity of Ado1.0 for adenosine and found that several structurally similar derivatives of Ado such as ADP, ATP, inosine, and adenine produced either a weak or undetectable response (Fig. 1H and fig. S2B).

Next, we examined whether Ado1.0 expression affects cellular physiology by measuring whether the sensor can activate downstream Gs protein–dependent signaling pathways. Using a luciferase complementation assay, we found that Ado1.0 has virtually no downstream Gs coupling; as a positive control, cells expressing A_2A_Rs produced robust downstream signaling (Fig. 1I, left and fig. S1E, middle). Similarly, Ado1.0 had no downstream cAMP activation induced by the A_2A_R agonist HENECA (Fig. 1I, right and fig. S1E, right). We also examined whether the Ado1.0 sensor can be internalized by applying 10 μM Ado to Ado1.0-expressing cells for 2 hours and found no significant decrease in fluorescence (Fig. 1J and fig. S2D), suggesting that the sensor remained at the cell membrane and does not undergo desensitization. Finally, we found no difference in either field stimulation–evoked Ca^2+^ signaling (fig. S3A-C and E) or K^+^-evoked glutamate release (fig. S3D and E) between Ado1.0-expressing neurons and non-transfected neurons, suggesting that expressing the Ado1.0 receptor did not measurably alter Ca^2+^ signaling or neurotransmitter release. Taken together, these results confirm that the Ado1.0 sensor is suitable for the use in both neurons and astrocytes, providing a sensitive and specific fluorescence response while sparing endogenous signaling.

### GRAB_Ado_ can monitor Ado dynamics *in vivo* during seizure activity and hypoxia in mice

Next, we validated the Ado1.0 sensor by monitoring Ado dynamics in freely moving mice. First, we examined the performance of the Ado1.0 sensor in a seizure model, in which kainic acid (KA) is injected into the hippocampus to induce seizures and increase Ado concentration in the hippocampus (*11*). We virally expressed Ado1.0 or the mutant Ado1.0mut in the CA1 region of the hippocampus and recorded the fluorescence signals using fiber photometry (Fig. 2A), and observed a large increase in fluorescence following intra-hippocampal KA injection in mice expressing Ado1.0, but not in mice expressing Ado1.0mut (Fig. 2B and C). Interestingly, the increase in Ado1.0 fluorescence signal was time-locked to the termination of each epileptic event measured using local field potential (LFP) recording, consistent with the notion that Ado signaling plays a role in terminating seizure activity (*12*). Next, we examined the performance of Ado1.0 during acute hypoxia (Fig. 2D), which has been shown to increase Ado concentrations (*13*). As expected, we found that inducing acute hypoxia in mice by N_2_ inhalation led to a rapid increase in Ado1.0 fluorescence in several brain regions, including the medial prefrontal cortex (mPFC) and nucleus accumbens (NAc) (Fig. 2E-G). Together, these results confirm that the Ado1.0 sensor can reliably report changes in Ado concentration in free moving mice under pathological conditions. Under physiological conditions, Ado1.0 is also capable of reporting rapid changes in Ado dynamics measured during natural sleep-wake cycles (*14*).

**Fig. 2.**
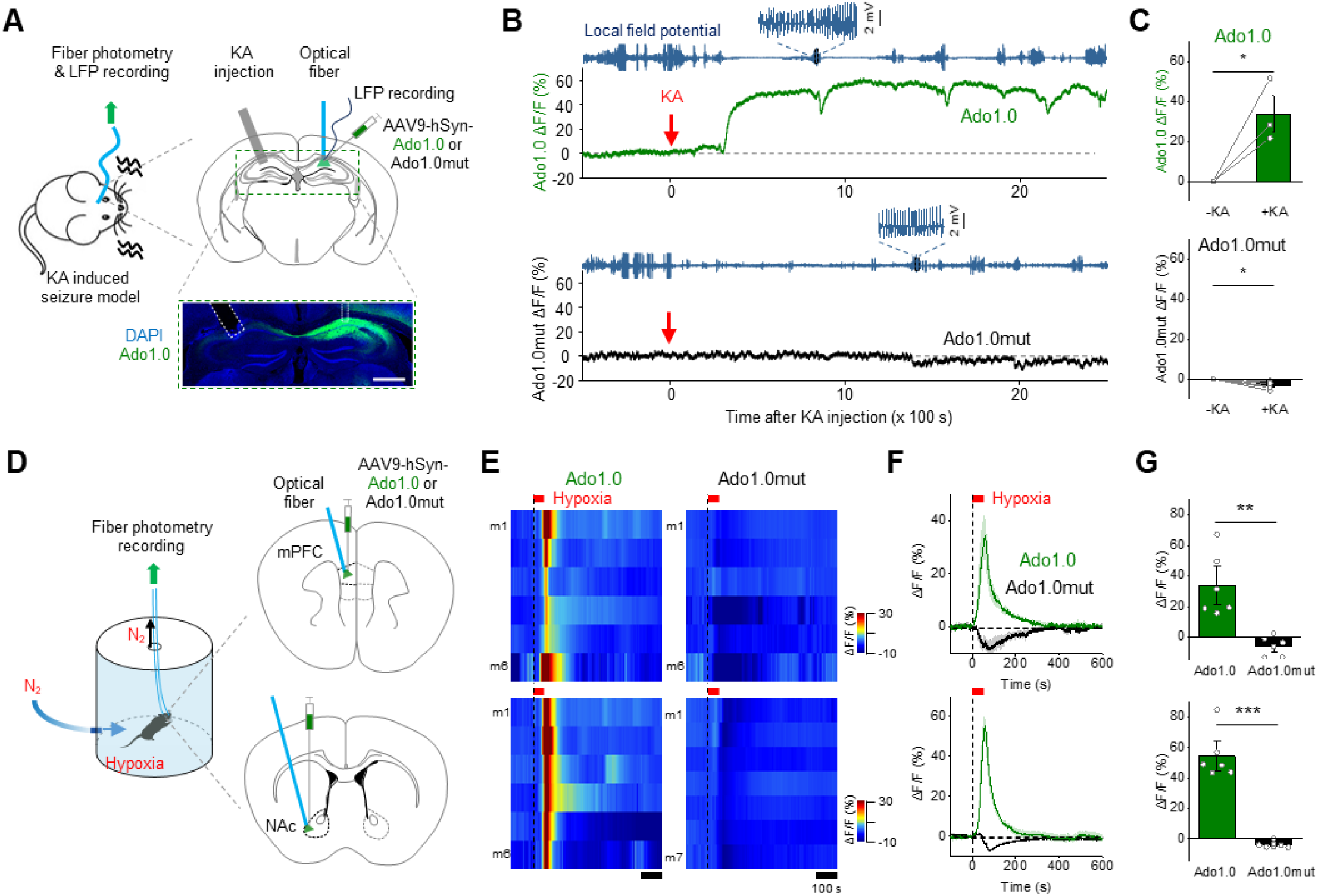
GRAB_Ado_ can monitor Ado dynamics *in vivo* during seizure activity and hypoxia in mice. **(A)** Schematic diagram depicting the strategy for using fiber photometry to record local field potentials (LFPs) and Ado1.0 or Ado1.0mut fluorescence in the hippocampus during seizure activity induced by kainic acid (KA) injected through a cannula. Shown below is an example of Ado1.0 expression in the hippocampus; the nuclei were counterstained with DAPI (scale bar, 1 mm). **(B and C)** Exemplar traces **(B)** and summary **(C)** of Ado1.0 and Ado1.0mut ΔF/F_0_; where indicated, KA was injected to induce epileptiform activity measured using LFP recording; n = 3-4 mice. **(D)** Schematic diagram depicting the strategy for using fiber photometry to record Ado1.0 or Ado1.0mut fluorescence in the medial prefrontal cortex (mPFC) and nucleus accumbens (NAc). Hypoxia was induced by infusing N_2_ into the chamber. **(E-G)** Heat maps **(E)**, averaged traces **(F)**, and summary **(G)** of Ado1.0 and Ado1.0mut ΔF/F_0_ measured in the mPFC (upper panels) and NAc (bottom panels). Where indicated, N_2_ was infused into the chamber to induce acute hypoxia; n ≥ 6 mice per group.

### GRAB_Ado_ reveals activity-induced Ado release from hippocampal neurons

A fundamental question in Ado signaling is the precise mechanism by which Ado is released in the brain. At the cellular level, Ado may be released from neurons (*15*) and/or from adjacent, non-neuronal cells such as astrocytes (*16*). At the molecular level, Ado may be produced by degradation of extracellular ATP released by various cell types (*17*) via the CD39-CD73 enzyme cascade (*18*). In addition, Ado may be released directly from neurons either via nucleoside transporters (*15*) or by vesicular exocytosis (*19*). To examine the cellular and molecular mechanisms that underlie the Ado release, we combined the use of Ado1.0 sensor with genetic and pharmacological experiments.

We first examined whether Ado release can be induced and measured by activating neurons and/or astrocytes. In cultured hippocampal neurons expressing Ado1.0, either electrical field stimuli (100 pulses at 30 Hz) or application of high extracellular K^+^ (75 mM) evoked robust and long-lasting fluorescence increases in both the soma and neuronal processes (Fig. 3A). Moreover, the stimulus-evoked increase in Ado1.0 fluorescence was blocked by the A_2A_R antagonist SCH-58261 (fig. S4A), suggesting specificity. Notably, we found a small transient decrease in Ado1.0 fluorescence in the presence of SCH-58261, likely caused by the neuronal activity-associated intracellular pH changes as previously reported (*20*). Ado1.0mut-expressing neurons showed similar transient fluorescence reduction (fig. S4), providing support to this speculation.

**Fig. 3.**
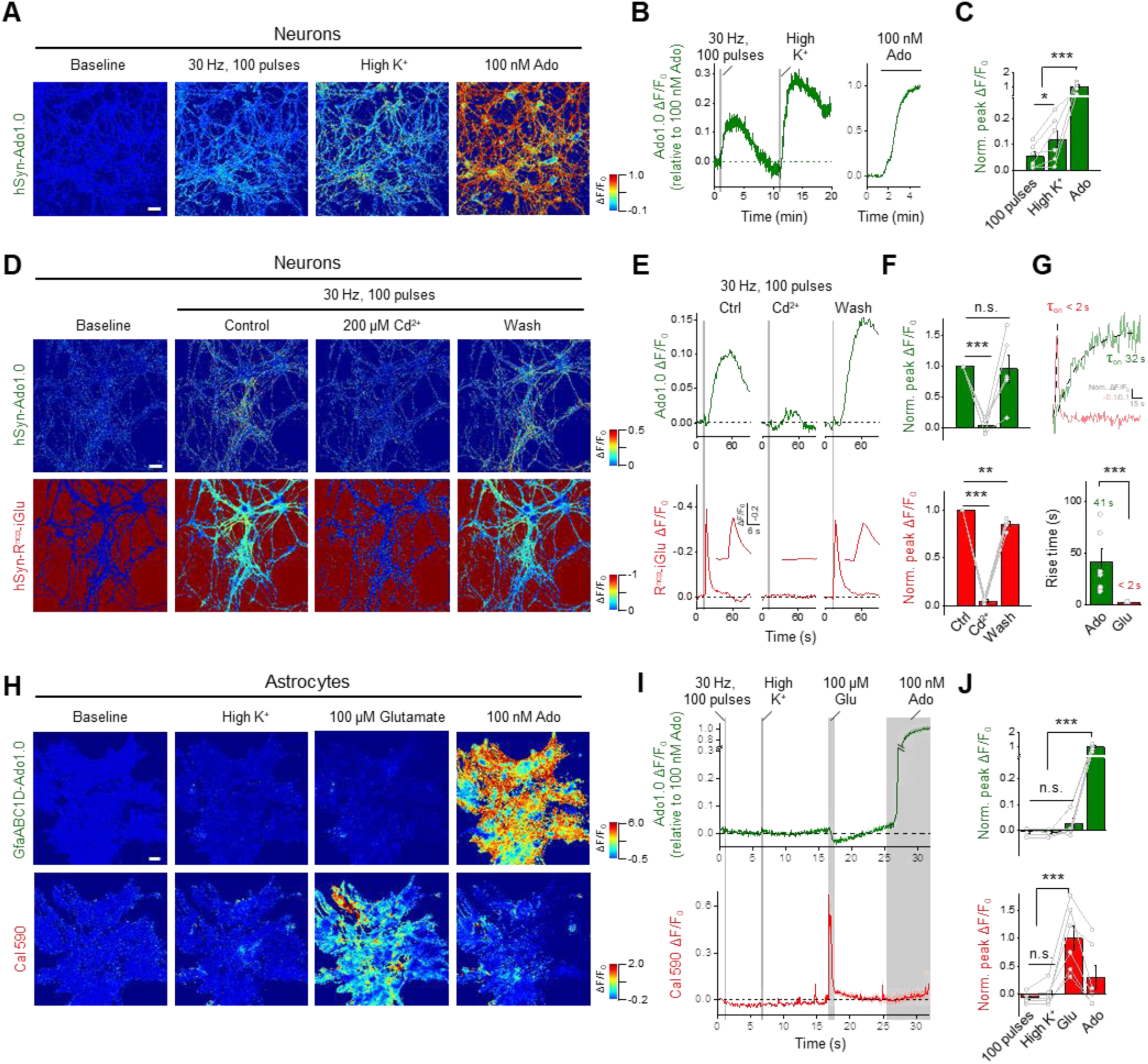
GRAB_Ado_ reveals activity-dependent Ado release in hippocampal neurons. **(A-C)** Ado1.0-expressing cultured hippocampal neurons were stimulated with 100 field stimuli at 30 Hz, high K^+^, or 100 nM Ado. Exemplar pseudocolor images **(A)**, example traces **(B)**, and summary data **(C)** are shown; in this and subsequent panels, the scale bar represents 30 μm; n = 4-6 coverslips each. **(D-F)** Dual-color imaging of Ado1.0 (upper panels) and R^ncp^-iGlu (bottom panels) in response to 100 field stimuli at 30 Hz applied before (control), during, and after (wash) bath application of 200 μM Cd^2+^; n = 4-5 coverslips. **(G)** Exemplar traces (top) and summary data (bottom) showing the kinetics of Ado1.0 and R^ncp^-iGlu ΔF/F_0_; n = 4-6 coverslips. **(H-J)** Dual-color imaging of Ado1.0 (upper panels) and Calbryte 590 (Cal 590, bottom panels) in response to 100 field stimuli at 30 Hz, high K^+^, 100 μM glutamate (Glu), or 100 nM Ado; n = 6 coverslips each.

We next examined whether the neuronal activity–dependent Ado1.0 signal requires Ca^2+^ signaling by applying cadmium (Cd^2+^), a non-selective blocker of voltage-gated Ca^2+^ channels (VGCCs). We simultaneously imaged glutamate (Glu) and Ado by co-expressing the red fluorescent glutamate sensor R^ncp^-iGluSnFR (R^ncp^-iGlu) (*21*) and the green fluorescent Ado1.0 sensor in cultured hippocampal neurons. As expected, stimulation-evoked Glu release was blocked by Cd^2+^ (Fig. 3D-F). Moreover, Cd^2+^ also blocked Ado1.0 responses (Fig. 3D-F), suggesting that Ca^2+^ influx through VGCCs is required for electrical or high K^+^ stimulation-induced Ado release. Next, we determined that the rise-time constants were 41 s and <2 s, respectively, for stimulation-induced Ado and Glu release (Fig. 3G). The slow on-kinetics of the Ado signal was likely due to the slow release of Ado but not the sensor’s intrinsic kinetics, given that Ado1.0 can rapidly respond to extracellular Ado with 68 ms kinetics (Fig. 1C). When we expressed Ado1.0 in cultured hippocampal astrocytes, none of the stimuli tested—including electrical field stimulation (30 Hz, 100 pulses), high K^+^, Glu (100 μM), bradykinin (1 μM) and thrombin (30 nM) application— induced significant fluorescence signals, although Glu, bradykinin, and thrombin evoked robust intracellular Ca^2+^ responses in the astrocytes (Fig. 3H-J and fig. S5). Taken together, these results suggest that neuronal activation is the principal source of Ado release in cultured hippocampal cells.

### Activity-dependent release of Ado depends on nucleoside transporters

The relatively slow release of Ado in activated neurons suggests that this process is not likely to be mediated by synaptic vesicle fusion, that is known to be rapid process. To test this, we expressed the tetanus toxin light chain (TeNT), which cleaves synaptobrevin and prevents exocytosis (*22, 23*). As expected, neurons expressing TeNT were abolished of stimulation-evoked Glu release; in contrast, Ado release was not affected (Fig. 4A), suggesting that it does not require vesicle fusion. We next examined whether the Ado1.0 signal is mediated by degradation of extracellular ATP by either blocking CD39 or knocking out CD73, two key ectoenzymes for the ATP/ADP to Ado conversion (Fig. 4B1). We found no effects on stimulation-induced Ado1.0 signals, suggesting that extracellular ATP does not contribute to activity-dependent extracellular Ado increase in cultured hippocampal neurons (Fig. 4B). Lastly, we examined whether equilibrative nucleoside transporters (ENTs) mediate Ado release (Fig. 4C1). We found that application of the ENT inhibitors NBTI (5 μM) and dipyridamole (DIPY, 10 μM) reversibly inhibited high K^+^-induced Ado release but not Glu release (Fig. 4C and fig. S6), indicating that ENT activity plays a key role in activity-induced Ado release in hippocampal neurons.

**Fig. 4.**
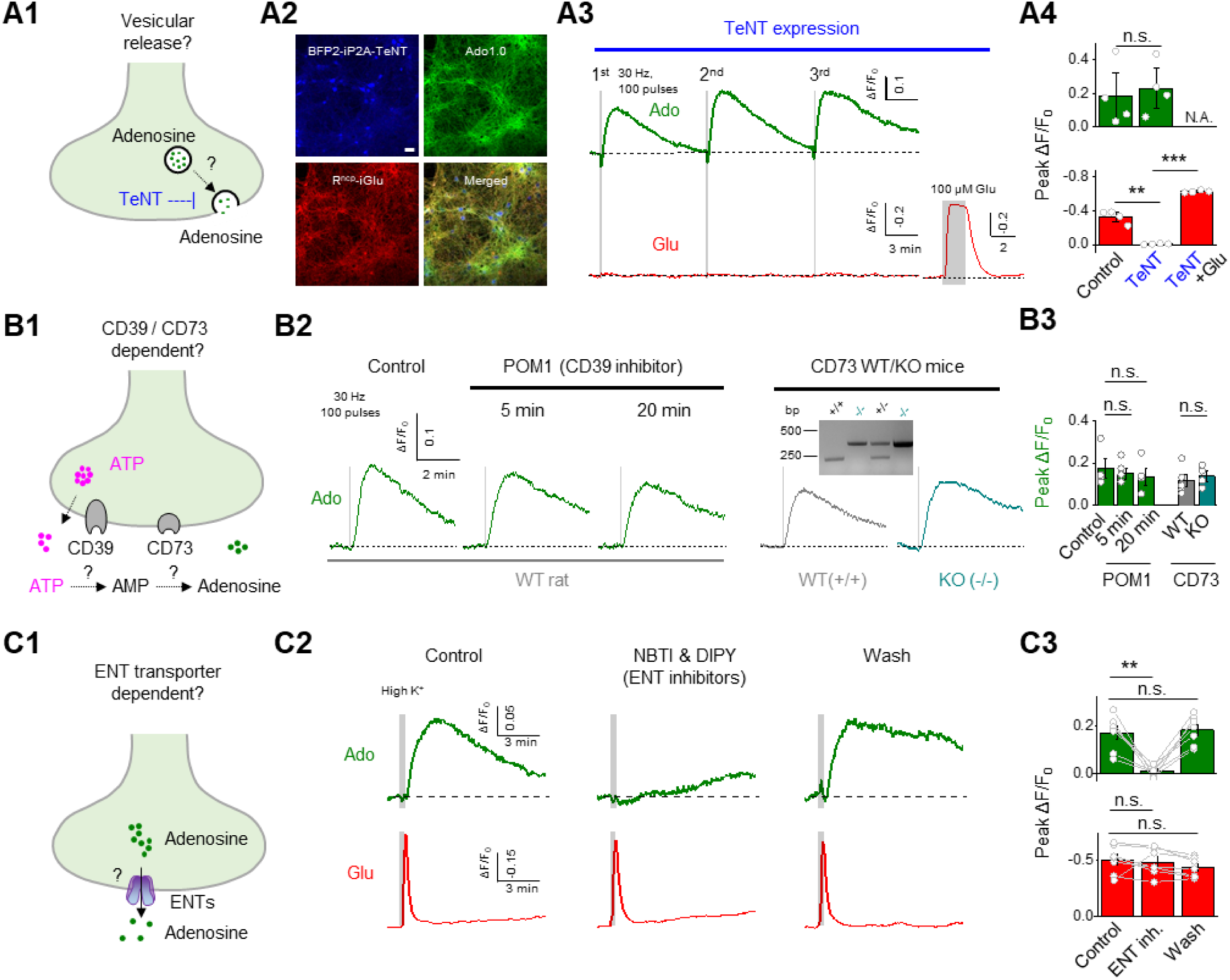
Activity-dependent Ado release is mediated by ENT transporters. **(A)** Blocking synaptic vesicle release with tetanus neurotoxin (TeNT) abolishes glutamate release, but not Ado release. **(A1)** Schematic drawing depicting the release of neurotransmitter-containing synaptic vesicles, which is blocked by TeNT. **(A2)** Representative confocal micrographs showing hippocampal neurons expressing TeNT (blue), Ado1.0 (green), and R^ncp^-iGlu (red). **(A3 and A4)** Averaged traces **(A3)** and group summary **(A4)** of Ado1.0 (upper panels, green) and R^ncp^-iGlu (bottom panels, red) ΔF/F_0_ in response to field stimuli (30 Hz, 100 pulses) with or without TeNT expression (n = 4 coverslips per group); the data in the control group are reproduced from fig. S6B. **(B)** Blocking CD39 or knocking out CD73 does not affect activity-dependent Ado release. **(B1)** Schematic drawing depicting the production of Ado from ATP via the CD39 and CD73 enzymes. **(B2 and B3)** Averaged traces **(B2)** and group summary **(B3)** of Ado1.0 ΔF/F_0_ in response to field stimuli (30 Hz, 100 pulses) under control conditions, in the presence of the CD39 inhibitor POM1 (10 μM), and in CD73 knockout neurons; n = 4-5 coverslips per group. The inset in **(B2)** shows example PCR genotyping results of CD73 KO mice. **(C)** Inhibiting ENT blocks activity-dependent Ado release, but not glutamate release. **(C1)** Schematic drawing depicting the release of Ado via equilibrative nucleoside transporters (ENTs). **(C2 and C3)** Averaged traces **(C2)** and group summary **(C3)** of Ado1.0 (upper panels, green) and R^ncp^-iGlu (bottom panels, red) ΔF/F_0_ in response to high K^+^ before (control), during, and after (wash) application of NBTI (5 μM) and dipyridamole (DIPY, 10 μM); n = 7 coverslips each.

### The Ca^2+^ source mediating activity-induced Ado release

Given the striking kinetics differences between neuronal glutamate- and Ado-release, it is possible that they may couple with distinct Ca^2+^ signaling, e.g. different subtypes of VGCCs. To test this, we used specific blockers of VGCC subtypes and simultaneously monitored glutamate and Ado release from the same neurons. We found that stimulation-induced Ado release was inhibited by nimodipine (Fig. 5A) and felodipine (fig. S7B), both of which block L-type VGCCs, the channel subtype primarily localized at the somatodendritic membrane in neurons (*24, 25*). Conversely, Ado release was largely unaffected by blocking P/Q-type and N-type VGCCs (Fig. 5A and fig. S7B), which are predominantly localized to presynaptic terminals and dictates Glu release (*26*). Taken together, these data suggest that the activity-induced Ado release occurs primarily at the somatodendritic region.

**Fig. 5.**
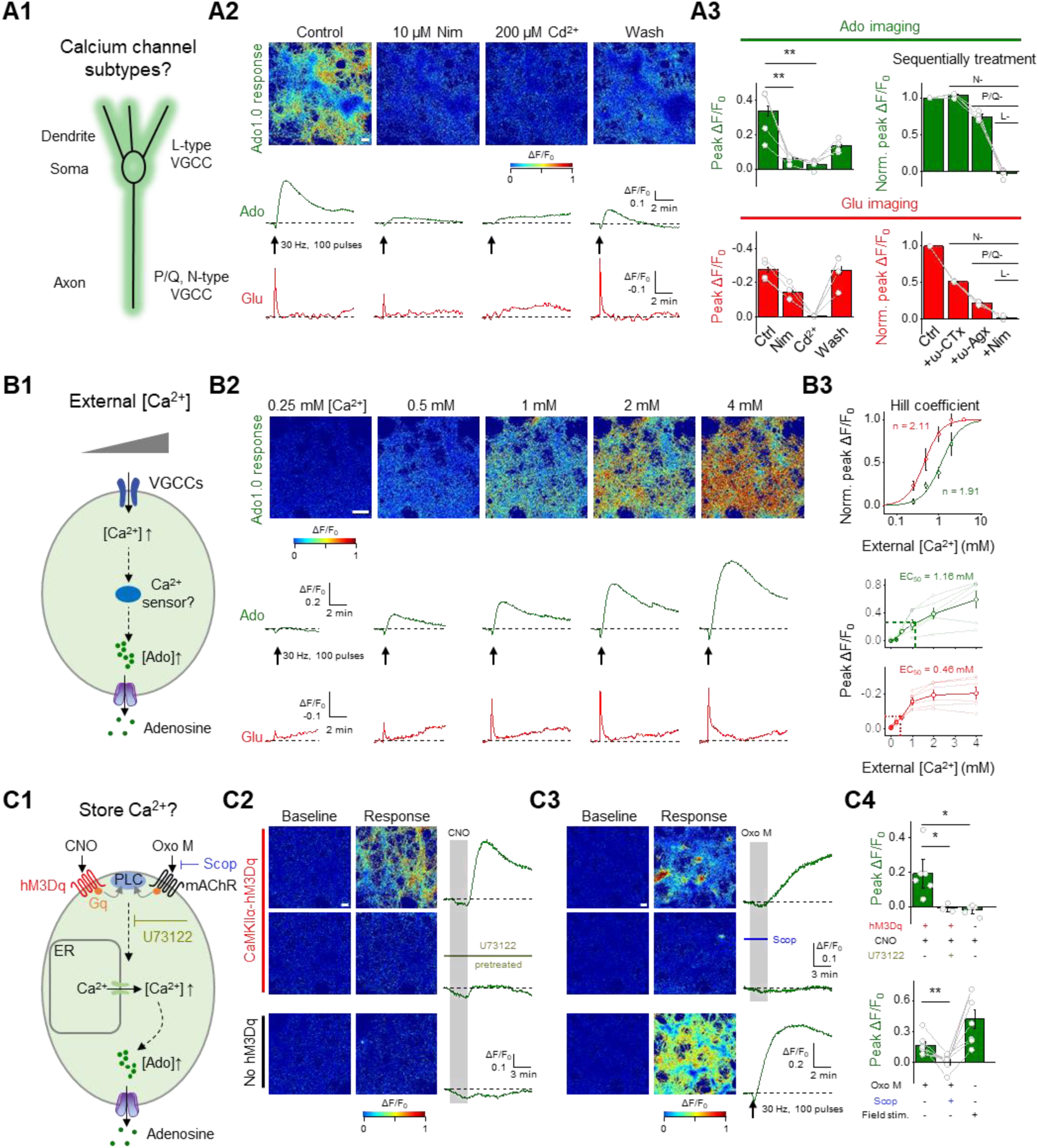
Dissecting the Ca^2+^ source that mediates activity-dependent Ado release. **(A)** Activity-dependent Ado release is blocked by inhibitors of L-type voltage-gated calcium channels (VGCCs). **(A1)** Schematic drawing depicting the expression of various VGCC subtypes in specific neuronal compartments. **(A2)** Pseudocolor images (upper panels) and exemplar traces (lower panels) of Ado1.0 and R^ncp^-iGlu ΔF/F_0_ in response to field stimuli (30 Hz, 100 pulses) applied before (control), during, and after (wash) application of nimodipine (Nim, 10 μM) or Cd^2+^ (200 μM); in this and subsequent panels, the scale bar represents 100 μm. **(A3)** Group summary of Ado1.0 (green) and R^ncp^-iGlu (red) ΔF/F_0_ in response to field stimuli, measured in the presence Cd^2+^ (200 μM) or Nim (10 μM), ω-Conotoxin-GVIA (ω-CTx, 1 μM), and ω-Agatoxin-IVA (ω-Agx, 0.3 μM) to block L-type, N-type, and P/Q-type VGCCs respectively; data shown in the left panels are from 4 independent coverslips, and the data shown in the right panels are from 4 ROIs in the same coverslip. **(B)** Ca^2+^ sensitivity differs between Ado release and Glu release. **(B1)** Schematic drawing depicting the experimental strategy. Dual-color imaging was used to image Ado and Glu release in various concentrations of extracellular Ca^2+^. **(B2)** Pseudocolor images (top) and averaged traces (bottom) of Ado1.0 and R^ncp^-iGlu ΔF/F_0_ in response to field stimuli (30 Hz, 100 pulses) in the indicated concentrations of extracellular Ca^2+^. **(B3)** Summary of peak Ado1.0 and R^ncp^-iGlu ΔF/F_0_ (n = 6 coverslips); the data in the top panel were normalized to the peak response measured in 4 mM Ca^2+^, and the data were fitted with a Hill equation (see main text). **(C)** Activation of Gq-coupled GPCRs induces Ado release. **(C1)** Schematic drawing depicting the experimental strategy. Clozapine N-oxide (CNO) and oxotremorine-M (Oxo-M) were used to activate recombinant hM3Dq-mCherry and endogenous Gq-coupled mAChRs, respectively; where indicated, the mAChR antagonist scopolamine (Scop) or the phospholipase C inhibitor U73122 was applied. **(C2)** Pseudocolor images (left) and example traces (right) of Ado1.0 ΔF/F_0_ in CaMKII1a-hM3Dq–expressing neurons in response to CNO (5 μM) in the absence (top) or presence (middle) of U73122 (10 μM); shown below are neurons that do not express CaMKII1a-hM3Dq. **(C3)** Pseudocolor images (left) and example traces (right) of Ado1.0 ΔF/F_0_ in response to Oxo-M (10 μM) in the absence (top) or presence (middle) of Scop (1 μM); as a positive control, separate neurons were stimulated with 100 pulses at 30 Hz (bottom). **(C4)** Summary of Ado1.0 ΔF/F_0_ measured under the indicated conditions; n = 3-7 coverslips.

To quantify the relative Ca^2+^ sensitivity of both Ado and Glu release in the same neurons, we measured the fluorescence responses of Ado1.0 and R^ncp^-iGlu as a function of external Ca^2+^ concentration. Ado1.0 and R^ncp^-iGlu fluorescence was imaged while applying electrical stimuli (100 pulses at 30 Hz) at increasing concentrations of extracellular Ca^2+^ (ranging from 0.25 to 4 mM, Fig. 5B). We then normalized the response measured at each Ca^2+^ concentration to the response measured at 4 mM Ca^2+^, and generated Ca^2+^-response curves (Fig. 5B3). To calculate the apparent affinity for extracellular Ca^2+^, the dose-response curve was fitted with the following Hill function: *(ΔF/F*_*0*_ *= 1 / (1*+ *(EC*_*50*_ */ [Ca*^*2*+^])^*n*^) as previously reported (*27, 28*), where ΔF/F_0_ is the normalized Ado1.0 or R^ncp^-iGlu response, EC_50_ is the apparent dissociation constant for extracellular Ca^2+^, [Ca^2+^] is the extracellular Ca^2+^ concentration, and *n* is the apparent cooperativity (Hill coefficient). Interestingly, we found that the calculated Ca^2+^ affinity was lower for Ado release than for Glu release, with apparent EC_50_ values of 1.16 μM and 0.46 μM, respectively; moreover, the cooperativity of Ado release was slightly lower than for Glu release, with Hill coefficient values of 1.91 and 2.11, respectively. These results indicate that activity-induced Ado release requires more Ca^2+^ than Glu release, suggesting that the Ca^2+^ sensor’s affinity differs between Ado and Glu release machinery.

Next, we tested whether triggering Ca^2+^ release from internal Ca^2+^ stores (by activating Gq-coupled GPCRs at the cell membrane) can induce Ado release (Fig. 5C1 and fig. S7A). We found that Ado release could be triggered by either activating the recombinant hM3Dq M3 muscarinic receptors (Gq) in CaMKIIα-positive hippocampal neurons using CNO, or by activating the endogenous metabotropic acetylcholine receptors (mAChRs; Gq) using Oxo-M. The release was blocked by pretreating the cells with the phospholipase inhibitor U73122 (10 μM) and the mAChR antagonist scopolamine (1 μM), respectively (Fig. 5C). Based on these results, we conclude that both depolarization-induced Ca^2+^ influx through L-type VGCCs and Ca^2+^ release from intracellular stores can drive Ado release from hippocampal neurons.

### Imaging somatodendritic Ado release

Given the dominant role of L-type VGCCs in supporting Ado release, it is reasonable to speculate that activity-dependent Ado release may occur preferentially at the somatodendritic membrane. To test this hypothesis, we imaged the Ado release in the hippocampus brain slices with well-defined anatomical projections. Unfortunately, although Ado1.0 was capable to report activity-induced Ado release in both somatodendritic and axonal compartments in cultured hippocampus neurons (Fig. 6A), we were unable to detect robust fluorescence increase in acute brains slices of the hippocampus (or mPFC, data not shown), possibly due to high basal levels of Ado present in acute brain slices (*29*) that saturated the high-affinity Ado1.0 sensor. Therefore, we generated a medium-affinity sensor for use in acute brain slices by introducing the M270H mutation in Ado1.0 (Ado1.0med), which had decreased affinity (Ado1.0med, EC_50_ 3.6 μM) and enhanced dynamic range (Ado1.0med, ΔF/F_0_ 350%) in response to Ado (fig. S8A). We found that Ado1.0med could readily report the activity-induced Ado release in acute brain slices containing the mPFC (fig. S8B-G).

**Fig. 6.**
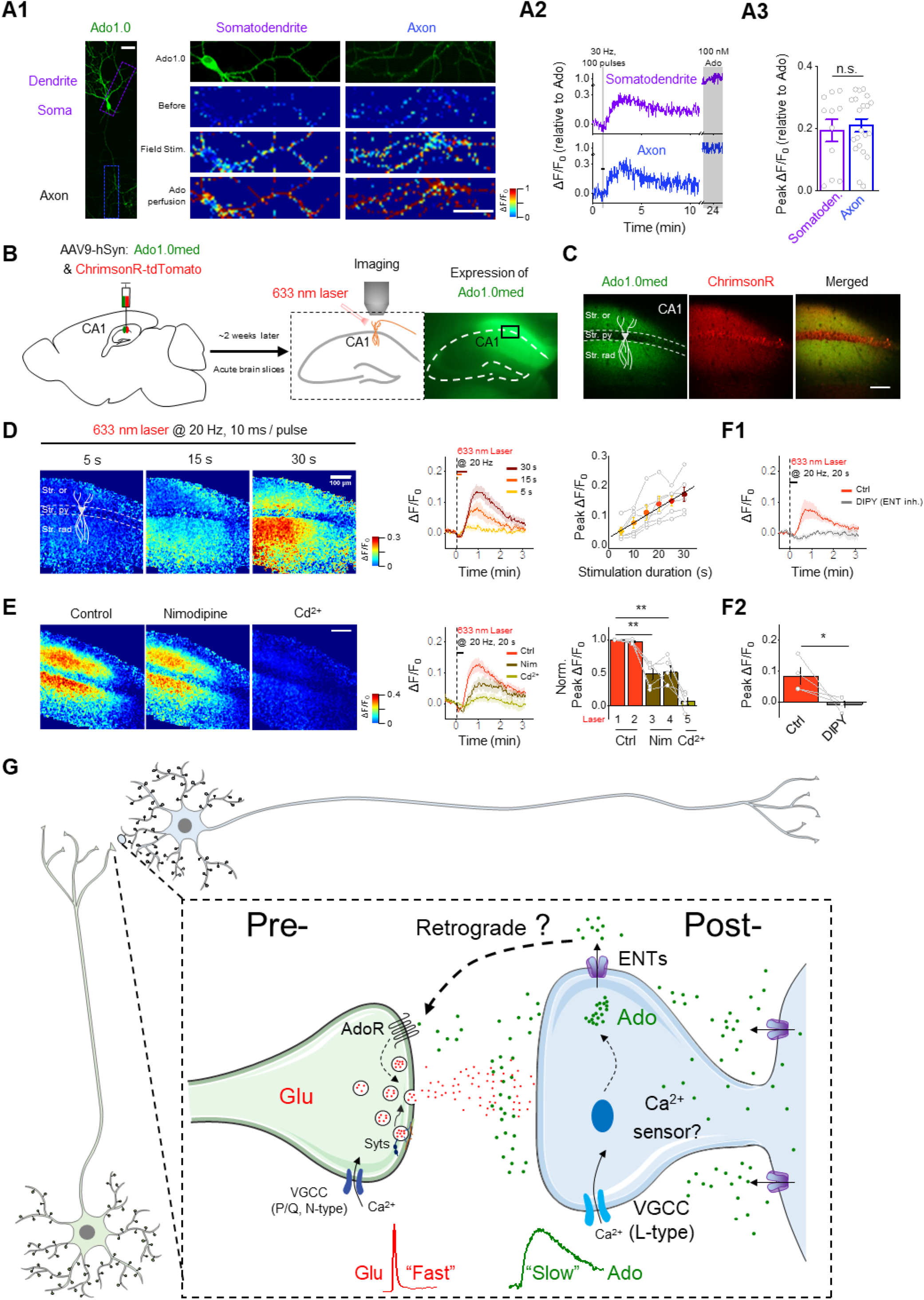
GRAB_Ado_ sensors reveal somatodendritic Ado release in hippocampal slices. **(A1)** Left panel: sparse expression of Ado1.0 in cultured hippocampal pyramidal neurons. Right panel: fluorescence images and pseudocolor images of Ado1.0 ΔF/F_0_ in the somatodendritic (purple box) and axonal (blue box) compartments in response to field stimuli (30 Hz, 100 pulses), high K^+^, or Ado (100 nM); scale bar, 50 μm. Example traces and summary data are shown in **(A2)** and **(A3)**, respectively; n = 11-21 ROIs from 4 coverslips per group. **(B)** Schematic illustration depicting the strategy used to image acute hippocampal brain slices prepared from mice expressing Ado1.0med and tdTomato-ChrimsonR in the CA1 region while using a 633-nm laser to activate the neurons. At the right is an image showing Ado1.0med fluorescence; the box shows the region magnified in **(C)**. **(C)** Magnified fluorescence images of the CA1 region showing Ado1.0med (green) and tdTomato-ChrimsonR (red) in CA1 regions (scale bar, 100 μm). The pyramidal cell layer (Str. py) is located between the stratum oriens (Str. ori) and stratum radiatum (Str. rad). **(D)** Pseudocolor images (left), averaged traces (middle), and group summary (right) of Ado1.0med ΔF/F_0_ in response to 633-nm laser pulses applied at 20 Hz for the indicated duration (scale bar, 100 μm); the solid line shown in the right panel a linear fit to the data; n = 6 slices from 4 mice. **(E)** Blocking L-type VGCCs inhibits optogenetically induced Ado release. Where indicated, nimodipine (Nim, 20 μM) and Cd^2+^ (100 μM) were applied (scale bar, 100 μm); n = 5 slices from 3 mice. **(F)** Blocking ENT transporters inhibits optogenetically induced Ado release. Where indicated, dipyridamole (DIPY, 20 μM) was used to block ENTs. Averaged traces **(F1)** and the group summary **(F2)** of Ado1.0med ΔF/F_0_ are shown; n = 4 slices from 3 mice. **(G)** Model depicting the novel mode of neuronal activity–dependent Ado release from hippocampal neurons. Ado is released slowly from the postsynaptic membrane via a vesicle-independent, ENT-dependent mechanism, serving as a putative retrograde signal to regulate presynaptic activity. This activity-dependent release of Ado requires L-type voltage-gated Ca^2+^ channels (VGCCs). In contrast, classic neurotransmitters such as glutamate (Glu) are released from presynaptic vesicles requires P/Q-type and N-type VGCCs. AdoR, adenosine receptor; ENTs, equilibrative nucleoside transporters; Syts, synaptotagmins.

Next, we expressed Ado1.0med in the CA3 region in acute hippocampal slices and recorded the fluorescence response in CA1 pyramidal neurons, which receive Schaffer collateral inputs from CA3 neurons (fig. S9A). Two-photon imaging confirmed the expression of Ado1.0med on CA3 axons innervating the CA1 region (fig. S9A). Increasing the intensity of electrical stimulation (20 Hz for 5 s) produced a progressively stronger Ado1.0med response (fig. S9B and C); moreover, the response was unaffected by co-application of the AMPA receptor antagonist NBQX and the NMDA receptor antagonist D-AP5 (fig. S9D).

We then asked whether Ado detected at Schaffer collateral terminals is directly released from dendrites of CA1 pyramidal neurons, where L-type VGCCs are enriched. To test this, we co-expressed the tdTomato-tagged light-activated ChrimsonR protein and Ado1.0med in hippocampal CA1 neurons, allowing us to use optogenetics to specifically activate CA1 neurons while simultaneously measuring cell-autonomous activation-evoked Ado release (Fig. 6B and C). Increasing numbers of 633-nm laser pulses applied at 20 Hz induced a progressively stronger fluorescence response in the soma, apical dendrites, and basal dendrites of CA1 pyramidal neurons (Fig. 6D), consistent with somatodendritic Ado release. Similar results were obtained using chemogenetic activation of CA1 pyramidal neurons, and—similar to the previous results—the response was unaffected by application of NBQX and D-AP5 (fig. S9E-H). Finally, consistent with the findings in cultured hippocampal neurons, inhibiting either L-type VGCCs or ENT transporters greatly attenuated the CA1 activity–induced release of Ado (Fig. 6E and F). Taken together, our results reveal the presence of a novel mode of Ado release from the somatodendritic compartment of hippocampal neurons (Fig. 6G).

## Discussion

Here, we report the development and characterization of genetically encoded fluorescent sensors GRAB_Ado_, which exhibits 60 nM EC_50_, 100 ms or less ON kinetics for adenosine (Ado). GRAB_Ado_ can readily report endogenous Ado dynamics during seizure and hypoxia with unprecedented sensitivity, stability and temporal resolution in living mice. Capitalizing on this unique tool, we unraveled pathways and mechanisms that regulate Ado release from neurons.

What is the source of extracellular Ado? We found that upon stimulation, Ado is predominantly released from neurons, but not from astrocytes, consistent with previous findings (*15*). Furthermore, we found the activity-dependent increase in Ado is due to direct release from the neuron, rather than indirectly from the degradation of extracellular ATP. Nevertheless, we cannot exclude the possibility that astrocytes can also release Ado or ATP under other conditions (*16, 30*). In this respect, our new GRAB_Ado_ sensors may serve as a valuable tool for examining this possibility in the future.

What are the mechanisms underlying Ado release? Using GRAB_Ado_ sensors combined with a spectrally distinct glutamate sensor, we found that the kinetics of Ado release was much slower than vesicular release of glutamate from the same neurons. Importantly, we found that the inhibition of ENT transporters, but not exocytosis machinery, significantly abolished the Ado release, without affecting glutamate release. Finally, in the same neuron, depolarization-induced Ado release required higher extracellular Ca^2+^ than glutamate release, suggesting different Ca^2+^ sensor affinity for Ado and glutamate. These data together reveal a unique routes and molecular pathways that control Ado release, distinct from the conventional vesicular- and SNARE-dependent pathway for classic neurotransmitters, such as glutamate. However, it is still unclear whether Ca^2+^ elevation is required for the generation of intracellular Ado, or the Ado transportation through ENTs, or both.

With respect to the site of Ado release from neurons, we found that the Ado release was very sensitive to blockers for L-type voltage-gated calcium channel (VGCC), a subtype known to primarily localized at the somatodendritic region. Moreover, loading an individual hippocampal CA1 pyramidal neuron with Ado was previously reported to selectively inhibit excitatory postsynaptic potentials, consistent with Ado release from the somatodendritic compartment (*31*). In acute hippocampal brain slices, using GRAB_Ado_ we directly visualized somatodendritic release of endogenous Ado from pyramidal neurons either by cell-autonomous optogenetic or chemogenetic activation of CA1 neurons. Given the abundance of presynaptic A_1_/A_2A_ receptors and the known presynaptic inhibition function of Ado in distinct brain regions (*31-33*), our work suggests that the somatodendritic release of Ado could serve as a retrograde modulator to feedback control presynaptic release.

Our results also indicate that the GRAB strategy can be used to develop sensors for measuring other purinergic transmitters (*34*). The availability of a comprehensive GRAB toolbox for purinergic transmitters, combined with robust emerging techniques such as optogenetics/chemogenetics and CRISPR/Cas9 will provide new insights into the role of purinergic transmission in both physiological and pathological conditions.

## Acknowledgments

We thank the members of the Li lab for suggestions and comments; we thank Drs. J-F Chen, M-M Poo, J-T Lv and Y-H Huang for critical reading of the manuscript.

## Author contributions

Y.L. conceived and supervised the project. Z.W., Huan W., Hao W, and Y.W. performed the experiments related to sensor development and optimization in cultured cells, with help from S.P., A.D., and M.J. Z.W. performed the experiments related to studying the mechanisms of Ado release in cultured neurons. Y.L., M.L., Z.W., Y.C., and A.D. designed and performed the experiments in acute brain slices. M.L., M.X., Z.Y., K.S. and W.P. designed and performed the *in vivo* mice experiments. All authors contributed to the data interpretation and data analysis. Y.L., Z.W., and M.X. wrote the manuscript with input from all other authors.

## Data and materials availability

All data necessary to assess the conclusions in this manuscript are provided in the manuscript and supplementary materials.

## Supplementary figure legends

**Fig. S1.**
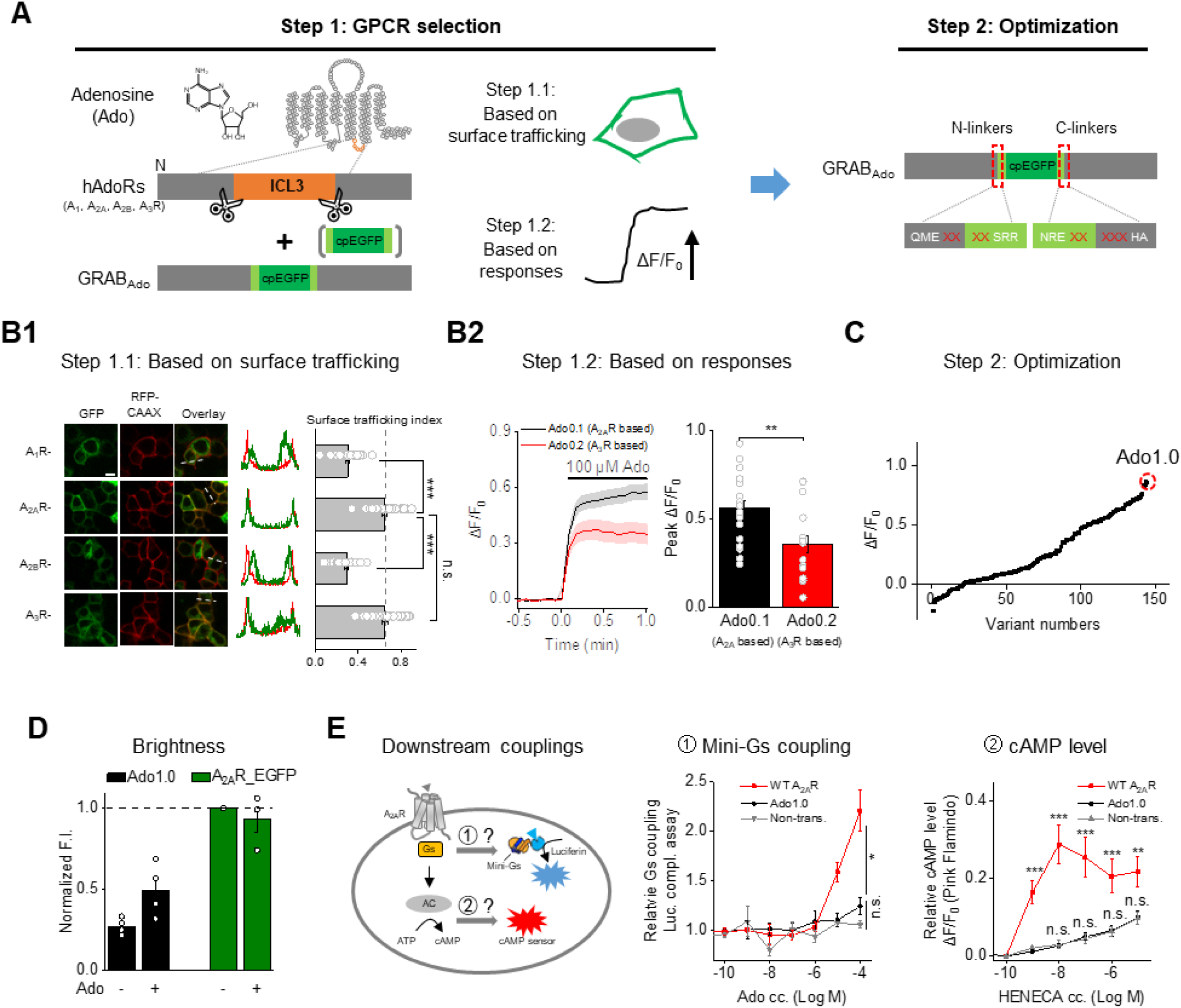
Design, optimization, and characterization of genetically encoded fluorescent Ado sensors in cultured cells (related to Fig. 1). **(A)** Schematic diagram depicting the strategy to screen the candidate GPCR scaffolds and optimize the adenosine (Ado) sensor. **(B)** Screening of the GPCR scaffold based on surface trafficking **(B1)** and the response to Ado **(B2). (B1)** Example images (left), line scans (middle), and summary data (right) of cell membrane trafficking of all four AdoRs (A_1_R, A_2A_R, A_2B_R, and A_3_R) with cpEGFP inserted (scale bar, 10 μm); n = 31 cells from 2-3 cultures per group. **(B2)** Summary of A_2A_R-based (Ado0.1) and A_3_R-based (Ado0.2) sensors in response to 100 μM Ado; n = 14-19 cells from 1-2 cultures. **(C)** Optimization of the N- and C-terminal linkers between A_2A_R and cpEGFP (indicated by dashed red boxes in panel **A**), yielding the most responsive Ado sensor, GRAB_Ado1.0_ (Ado1.0). Shown is the fluorescence change measured in response to 20 μM Ado, normalized to Ado1.0 (indicated with a dashed red circle). **(D)** Relative fluorescence intensity of Ado1.0 and an EGFP-tagged A_2A_R measured in the absence and presence of 100 μM Ado, normalized to control; n ≥ 3 coverslips with ≥ 10 cells per coverslip. **(E)** Ado1.0 does not signal to downstream Gs pathways (related to **Fig. 1I**). Left, schematic diagram depicting the strategy used to measure downstream Gs protein coupling. Shown at the right are a luciferase complementation assay to measure Gs protein coupling and the cAMP sensor PinkFlamindo to measure cAMP levels in non-transfected HEK293T or HeLa cells and cells expressing A_2A_R or Ado1.0 following application of the indicated concentrations of Ado or the A_2A_R agonist HENECA; n ≥ 3 independent experiments per group.

**Fig. S2.**
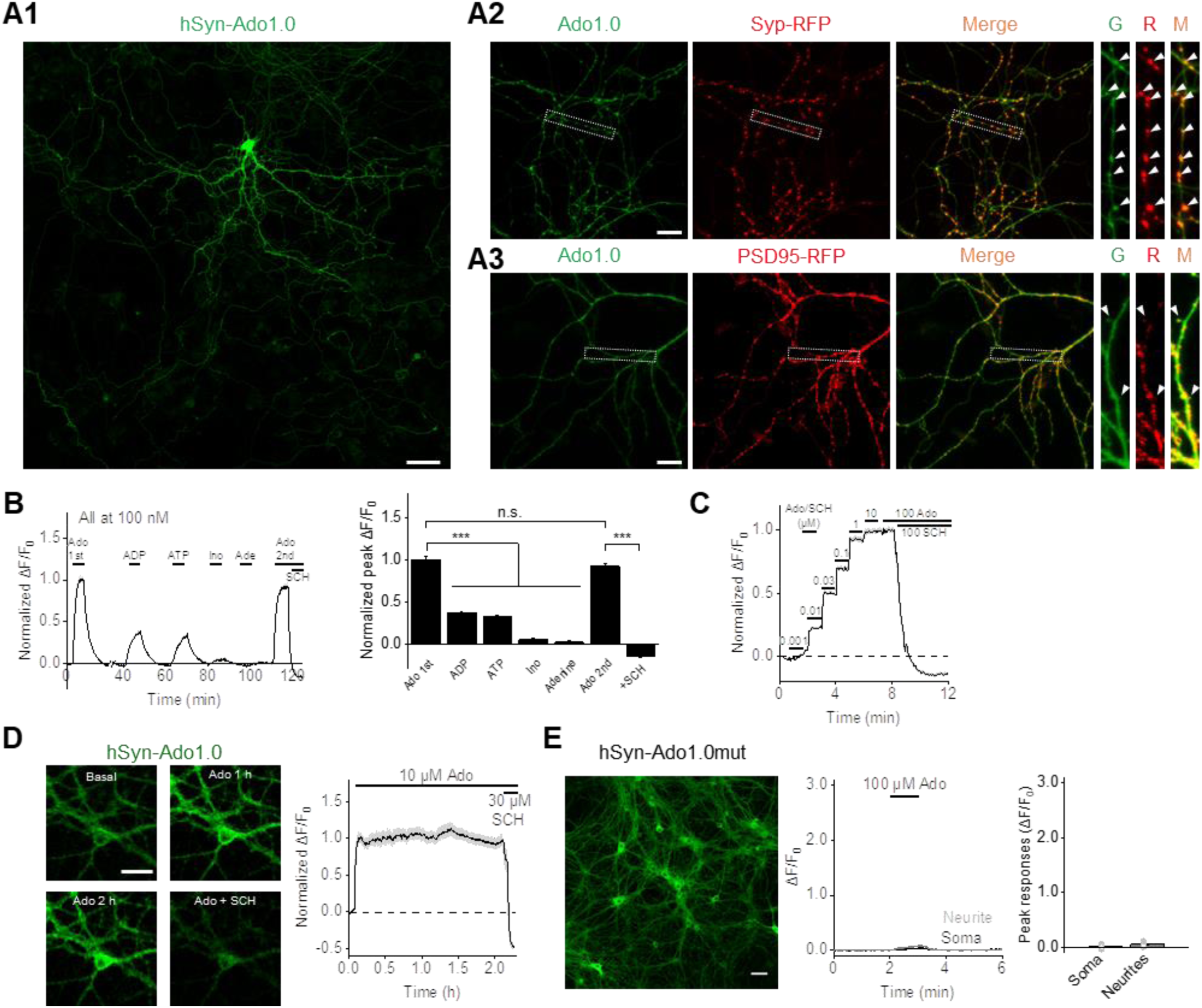
Characterization of GRAB_Ado1.0_ sensors in cultured neurons (related to Fig. 1). **(A)** Expression and localization of the Ado1.0 sensor (green) and subcellular markers (red) in the indicated subcellular compartments in cultured neurons. RFP-tagged synaptophysin (Syp-RFP) and PSD95 (PSD95-RFP) were co-expressed with Ado1.0 to label the presynaptic boutons and postsynaptic dendritic spines, respectively (indicated by arrowheads); scale bars, 50 μm **(A1)** and 20 μm **(A2 and A3)**. **(B)** Example recording and summary of normalized ΔF/F_0_ in Ado1.0-expressing neurons in response to the indicated compounds (each at 100 nM); n = 10 neurons from 1 coverslip. Ado, adenosine; ADP, adenosine diphosphate; ATP, adenosine triphosphate; Ino, inosine; Ade, adenine; SCH, SCH-58261. **(C)** Normalized ΔF/F_0_ in Ado1.0-expressing neurons in response to the indicated concentrations of Ado, followed by the A_2A_R antagonist SCH; the trace represents the average of 10 ROIs from a single coverslip. **(D)** Example fluorescence images and averaged ΔF/F_0_ trace of Ado1.0-expressing neurons induced by 10 μM Ado for 2 hours (scale bar, 30 μm); n = 28 neurons from 3 cultures (related to **Fig. 1J**). **(E)** Ado1.0mut-expressing neurons do not have a fluorescence response to 100 μM Ado. Shown are an example confocal GFP fluorescence image (left panel; scale bar, 50 μm), example time course of ΔF/F_0_ (middle panel), and summary of peak ΔF/F_0_ (right panel); n = 56 ROIs from 4 cultures.

**Fig. S3.**
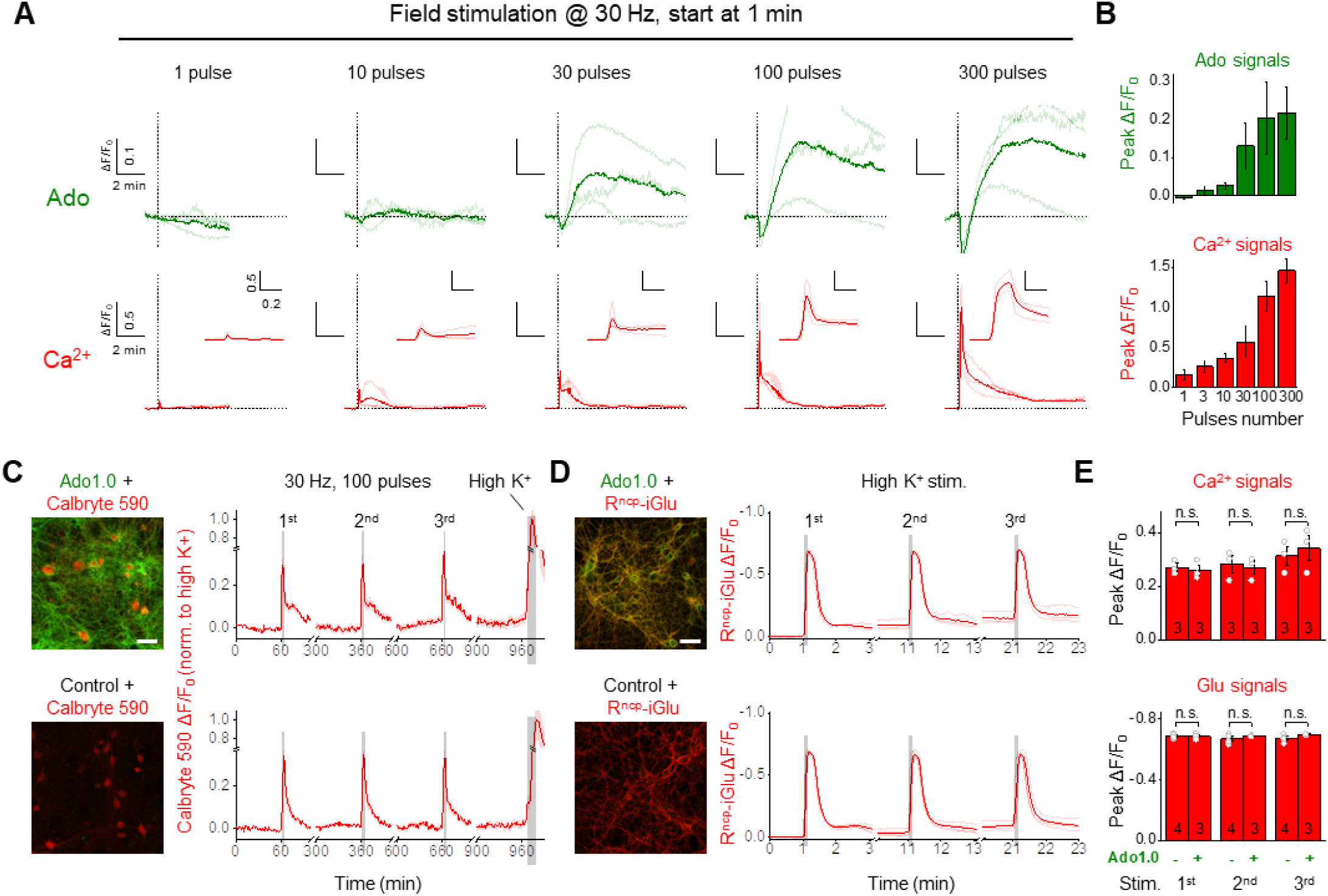
Expressing Ado1.0 does not affect Ca^2+^ signaling or glutamate release (related to Fig. 1). **(A and B)** Traces **(A)** and group summary **(B)** of Ado1.0 and the Ca^2+^ indicator Calbryte 590 ΔF/F_0_ in response to the indicated number of field stimuli applied at 30 Hz. In **(A)**, the thin lines represent individual cells, and the thick lines represent the average trace; insets in the lower row meant magnified view of 0.8-1.5 min of traces. **(C-E)** Calbryte 590 **(C)** and the genetically encoded glutamate sensor R^ncp^-iGlu **(D)** were used to measure Ca^2+^ and Glu release in Ado1.0-expressing neurons (upper panels) and control neurons (bottom panels). Confocal images (left panels; scale bars, 50 μm) and normalized raw traces (right panels) of Calbryte 590 and R^ncp^-iGlu ΔF/F_0_ in response to field stimuli (30 Hz, 100 pulses) or high K^+^ are shown, and the summary data are shown in **(E)**; n = 3-4 coverslips each.

**Fig. S4.**
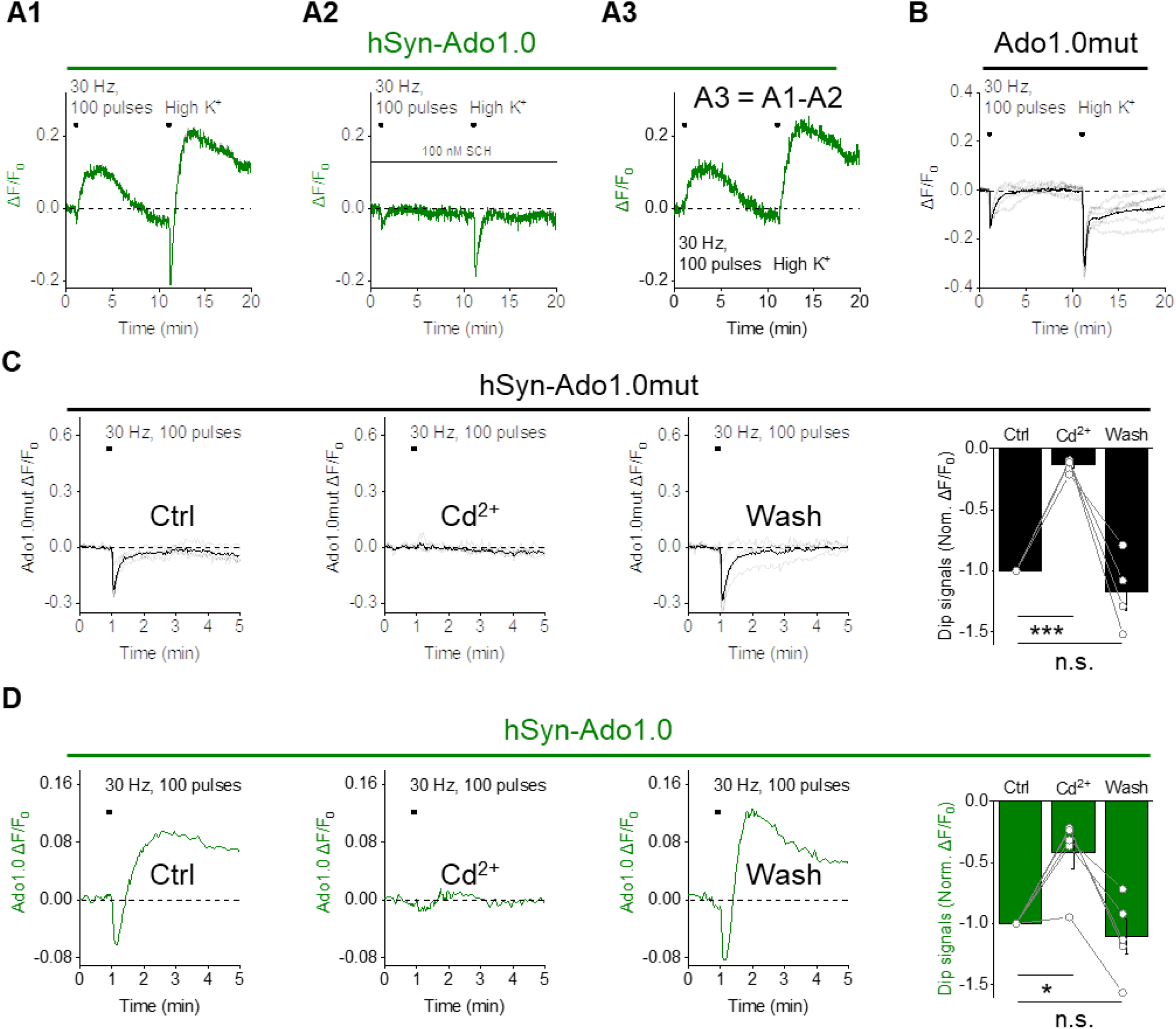
Specificity of stimulation-induced Ado1.0 ΔF/F_0_ in cultured hippocampal neurons (related to Fig. 3). **(A)** Exemplar traces of ΔF/F_0_ measured in Ado1.0-expressing cultured hippocampal neurons in response to 100 field stimuli delivered at 30 Hz and high K^+^ in the absence **(A1)** and presence **(A2)** of SCH-58261 (SCH, 100 nM). A3 shows the subtracted trace. **(B)** Traces showing Ado1.0mut ΔF/F_0_ in response to 100 field stimuli at 30 Hz and high K^+^ (n = 6 coverslips). **(C and D)** Ado1.0mut **(C)** and Ado1.0 **(D)** ΔF/F_0_ measured in response to 100 field stimuli delivered at 30 Hz before (Ctrl), during, and after (Wash) application of Cd^2+^ (200 μM). Summary data are shown at the right, normalized to control; n = 4-5 coverslips per group.

**Fig. S5.**
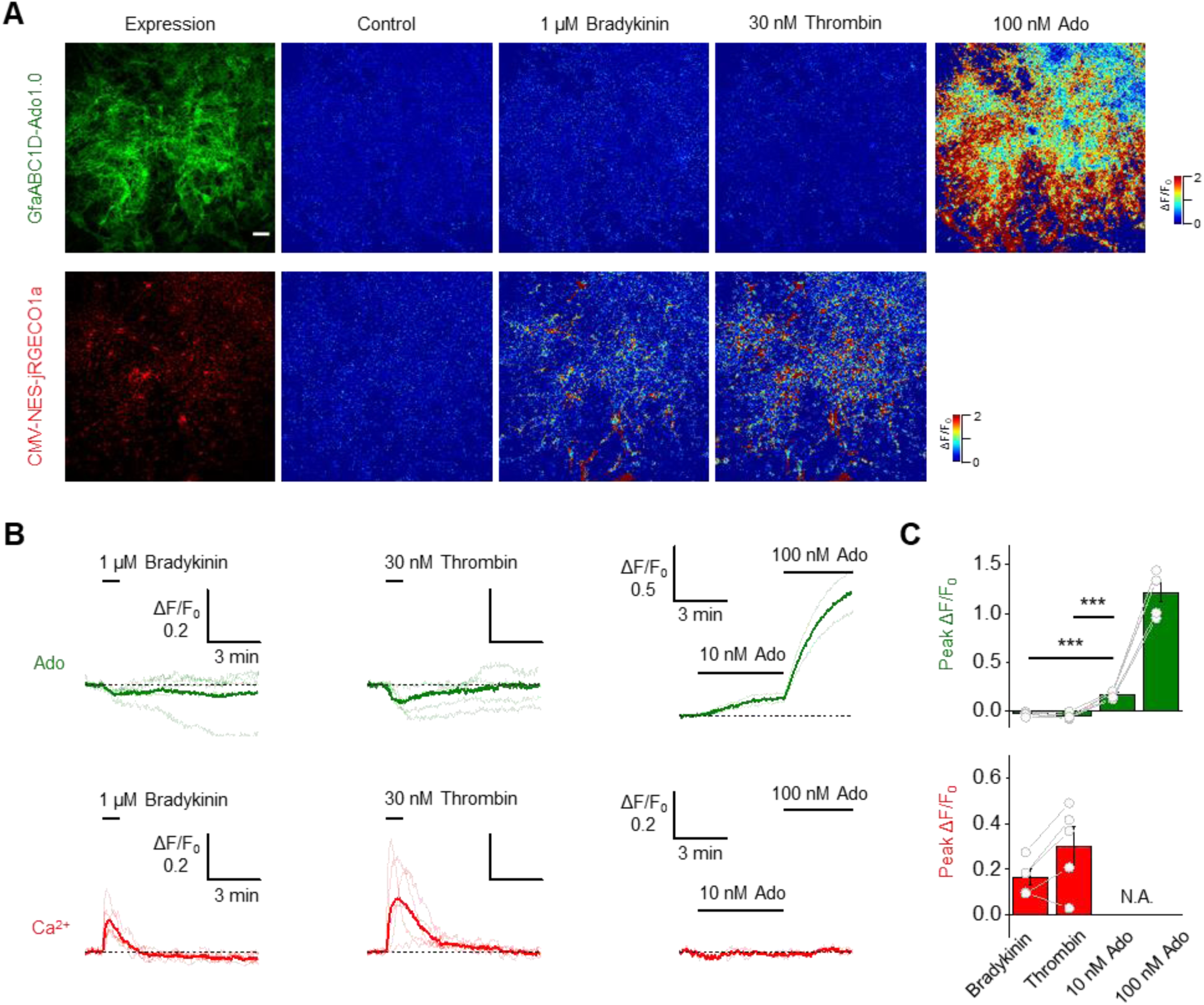
Agonists for Gq-coupled GPCRs did not induce Ado release from astrocytes (related to Fig. 3). **(A-C)** Dual-color imaging of Ado1.0 (upper panels) and NES-jRGECO1a (bottom panels) in response to 1 μM bradykinin, 30 nM thrombin, 10 nM Ado or 100 nM Ado; confocal or pseudocolor images **(A)**, traces **(B)** and group summary **(C)** are shown; scale bar, 100 μm; n = 5 coverslips each.

**Fig. S6.**
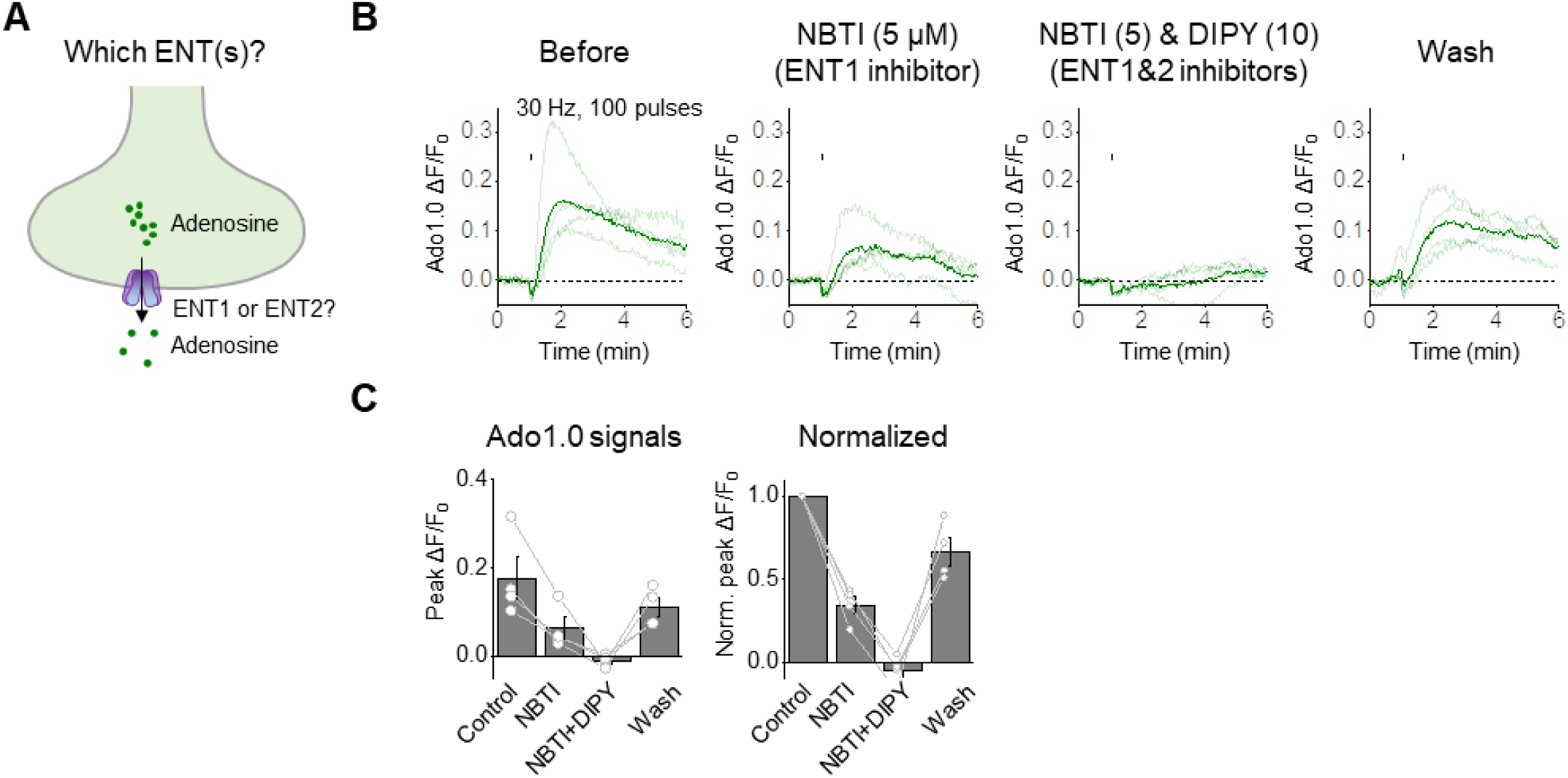
Both ENT1 and ENT2 contribute to activity-dependent Ado release (related to Fig. 4). **(A)** Schematic drawing depicting the putative mechanism by which equilibrative nucleoside transporters (ENTs) mediate Ado release in neurons. **(B and C)** Averaged traces **(B)** and group summary **(C)** of Ado1.0 ΔF/F_0_ in response to field stimuli (30 Hz, 100 pulses, indicated by black tick marks) before (control), during, and after (wash) application of the ENT1 blocker NBTI (5 μM) and/or the ENT1/2 blocker dipyridamole (DIPY, 10 μM); n = 4 coverslips.

**Fig. S7.**
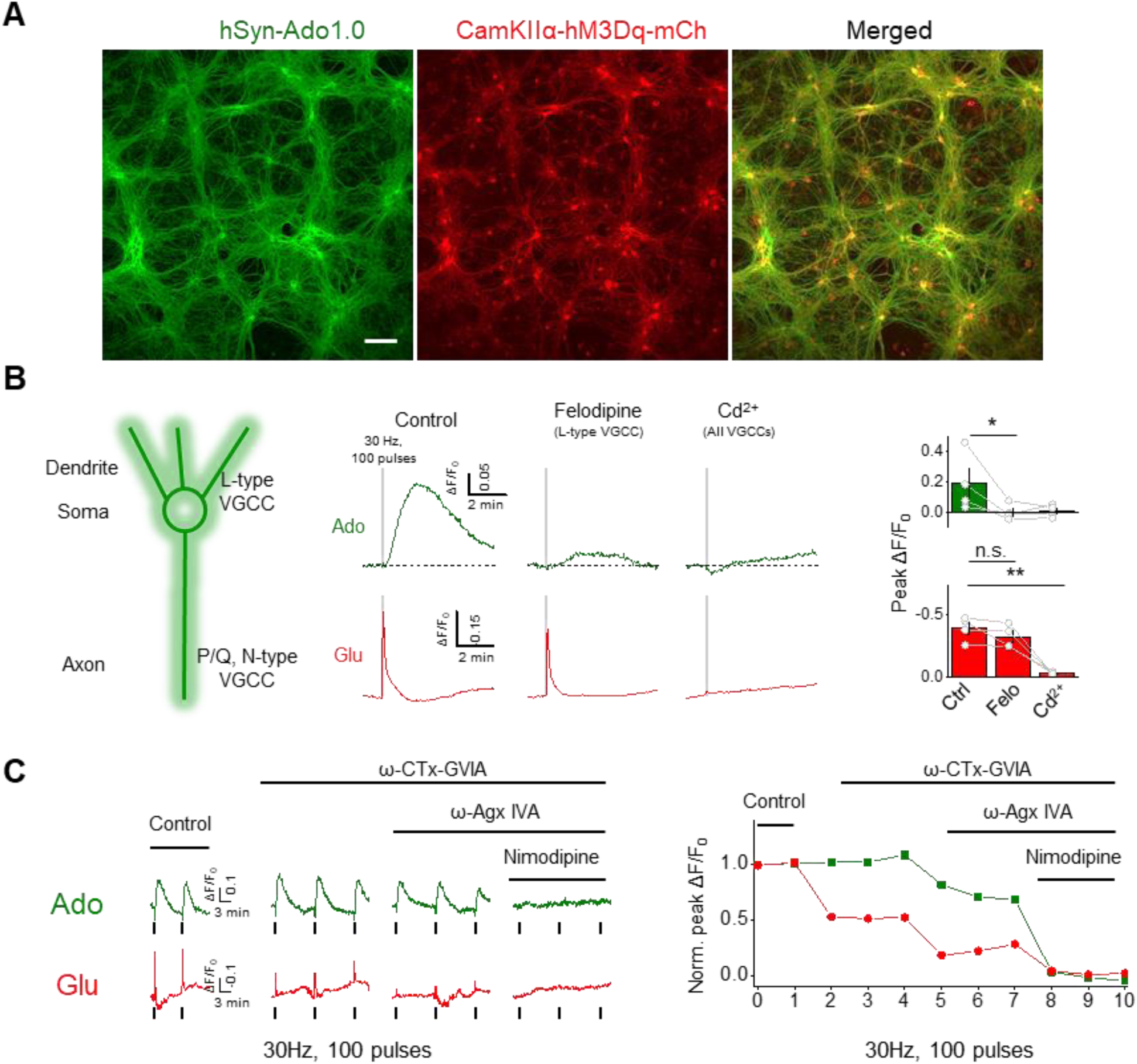
Dissecting the subcellular localization and Ca^2+^ source of activity-dependent Ado release in cultured neurons (related to Fig. 5). **(A)** Confocal images of cultured hippocampal neurons co-expressing Ado1.0 (green) and mCherry-tagged CaMKIIα-hM3Dq (red); scale bar, 100 μm. **(B)** Blocking L-type voltage-gated calcium channels (VGCCs) inhibits Ado release. Left panel: cartoon illustrating that L-type VGCCs are expressed primarily in the somatodendritic compartment, while P/Q-type and N-type VGCCs are expressed primarily in the axon. Middle panels: example traces of Ado1.0 and R^ncp^-iGlu ΔF/F_0_ in response to field stimuli (30 Hz, 100 pulses) delivered before (control), during, and after (wash) application of either the L-type VGCC blocker felodipine (Felo, 10 μM) or Cd^2+^ (200 μM). Right: group summary of Ado1.0 (top panel) and R^ncp^-iGlu (bottom panel) ΔF/F_0_ in response to field stimuli; n = 4 coverslips each. **(C)** Example traces and normalized peak Ado1.0 and R^ncp^-iGlu ΔF/F_0_ in response to field stimuli (30 Hz, 100 pulses) applied in the presence of ω-CTx-GVIA (1 μM), ω-Agx-IVA (0.3 μM), and nimodipine (10 μM) to block N-type, P/Q-type, and L-type VGCCs, respectively (the same coverslip was sequentially treated with the indicated blockers). Related to **Fig. 5B3**.

**Fig. S8.**
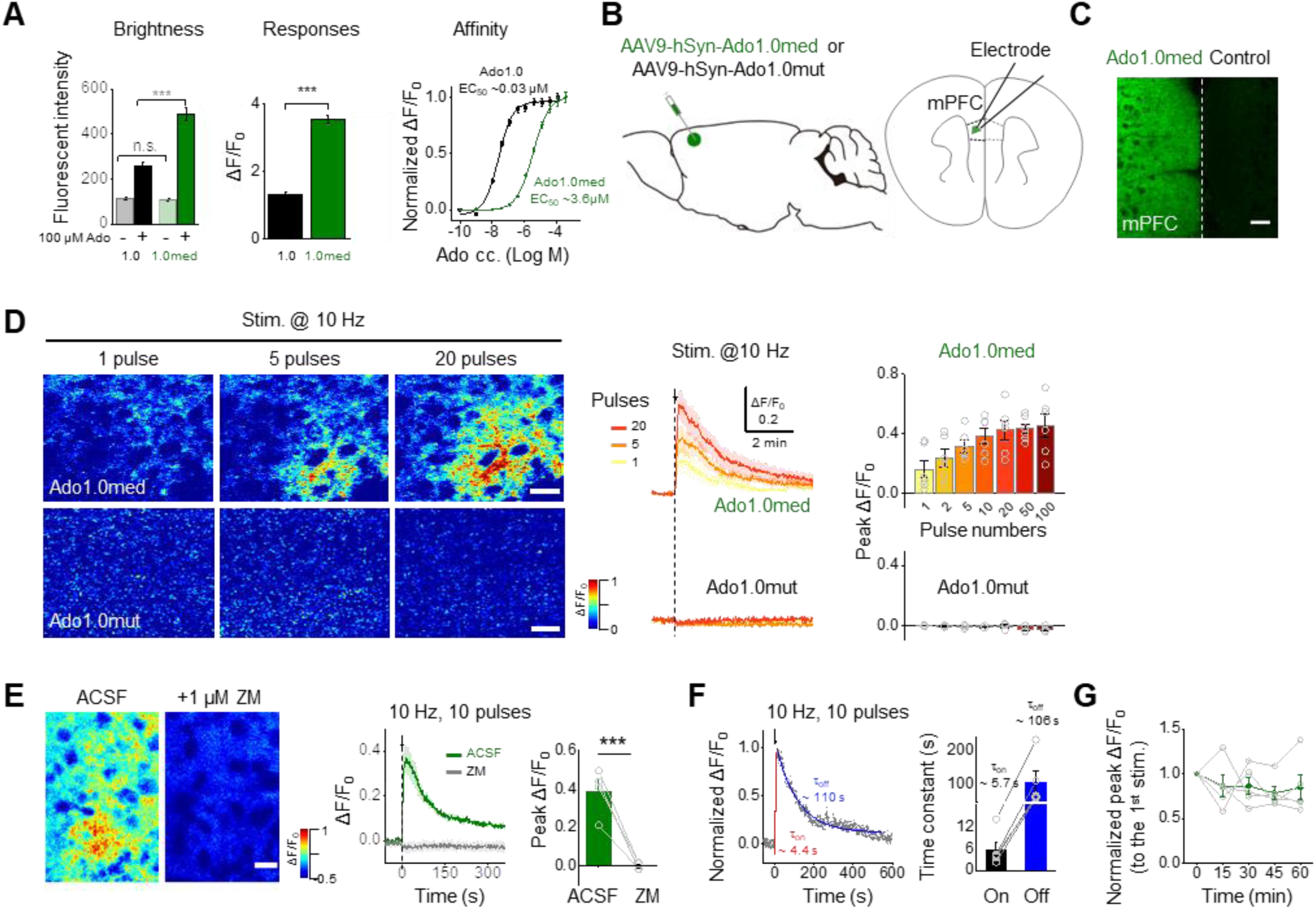
Characterization of Ado1.0med in cultured neurons and acute mPFC brain slices (related to Fig. 6). **(A)** Fluorescence intensity (left panel), peak ΔF/F_0_ (middle panel), and dose-response curves (right panel) for Ado1.0 and the medium-affinity Ado1.0med sensor in cultured neurons in response to adenosine (Ado); n = 20 ROIs from 2 cultures per group. **(B)** Schematic illustration depicting the expression of Ado1.0med or Ado1.0mut in the mouse mPFC, an acute brain slice containing the mPFC, and the placement of a bipolar stimulating electrode in the mPFC. **(C)** Fluorescence images showing the expression of Ado1.0med in the ipsilateral mPFC, with no expression in the contralateral mPFC (scale bar, 50 μm). **(D)** Pseudocolor images (left panels), example traces (middle panels), and group summary (right panels) of Ado1.0med ΔF/F_0_ in response to the indicated number of electrical stimuli delivered at 10 Hz (scale bar, 20 μm); n = 6 slices from 3 mice per group. **(E1)** Pseudocolor images (left panels), representative traces (middle panels), and group summary (right panels) of Ado1.0 ΔF/F_0_ in response to 10 electrical pulses at 10 Hz delivered in control solution (ACSF) or solution containing the A_2A_R antagonist ZM-241385 (ZM, 1 μM); scale bar, 20 μm. **(E2)** Time course (left) and summary (right) of Ado1.0med ΔF/F_0_ in response to the indicated number of electrical stimuli at 10 Hz; n = 4 slices from 1 mouse each. **(F)** Representative traces (left) and group summary (right) of normalized ΔF/F_0_ and kinetics (τ_on_ and τ_off_) in Ado1.0-expressing neurons in response to 10 electrical stimuli delivered at 10 Hz; n = 5 slices from 3 mice. **(G)** ΔF/F_0_ was measured in Ado1.0-expressing neurons in response to repeated trains of electrical stimuli applied every ∼15 min and normalized to the first train; n = 5 slices from 4 mice.

**Fig. S9.**
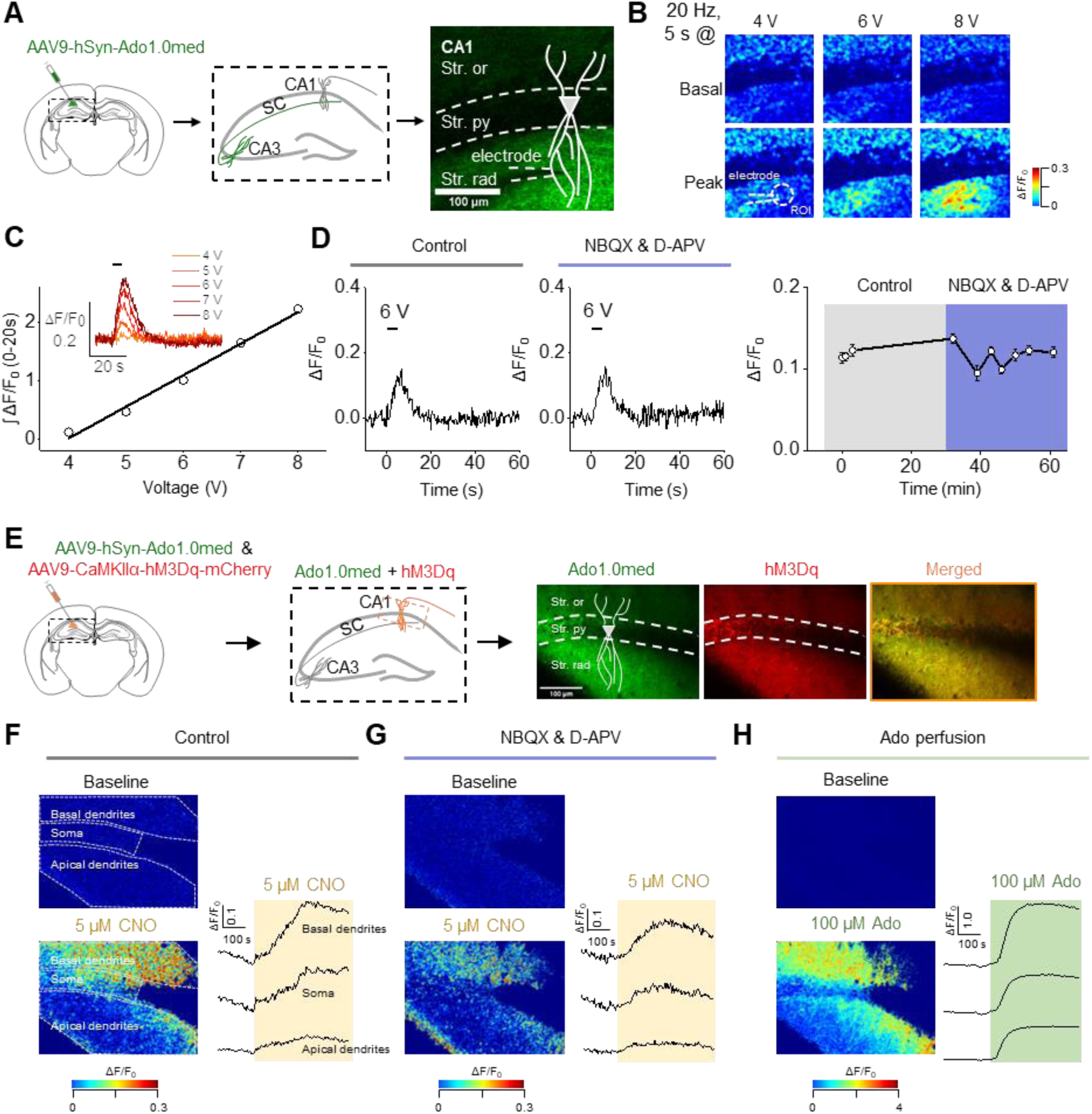
Imaging Ado release in acute hippocampal slices (related to Fig. 6). **(A)** Schematic drawing depicting the strategy for injecting AAV9-hSyn-Ado1.0med in the mouse hippocampal CA3 region (left) and expressing Ado1.0med in CA3 axons innervating the CA1 region; the image at the right shows the approximate location of the pyramidal cell body and neurites. **(B)** Pseudocolor images of Ado1.0med ΔF/F_0_ in response to electrical stimuli (20 Hz for 5 s, ranging in intensity from 4 V to 8 V). The white dashed circle indicates the region of interest (50 μm in diameter) used for quantification. **(C)** Raw traces (inset) and summary of Ado1.0med ΔF/F_0_ in response to electrical stimuli applied at the indicated voltages. **(D)** Raw traces (left panels) and summary (right panels) of Ado1.0med ΔF/F_0_ in response to electrical stimuli in control solution or solution containing NBQX (10 μM) and D-APV (50 μM). **(E)** Schematic drawing depicting the strategy for injecting AAV9-hSyn-Ado1.0med and AAV9-CaMKIIa-hM3Dq-mCherry in the mouse CA1 region (left), and fluorescence images showing the expression of Ado1.0med and hM3Dq-mCherry in CA1 neurons (right). **(F)** Pseudocolor images (left) and raw traces (right) of Ado1.0med ΔF/F_0_ before and after application of CNO (5 μM). The white dashed regions correspond to the ROIs used to measure ΔF/F_0_ in basal dendrites, the soma, and apical dendrites. **(G)** Same as **(F)**, except CNO (5 μM) was applied in the presence of NBQX (10 μM) and D-APV (50 μM). **(H)** Same as **(F)**, except Ado (100 μM) was applied. Note the different vertical scale (1.0) compared to panels **F** and **G** (0.1).

## Materials and Methods

### Molecular biology

Plasmids were generated using Gibson assembly (*1*). DNA fragments were generated using PCR amplification with primers (Tsingke) with ∼25-bp overlap, and all sequences were verified using Sanger sequencing. All cDNAs encoding the candidate GRAB_Ado_ sensors were cloned into the pDisplay vector (Invitrogen) with an upstream IgK leader sequence and a downstream IRES-mCherry-CAAX cassette (to label the cell membrane). The cDNAs encoding the Ado receptor subtypes A_1_R, A_2A_R, A_2B_R, and A_3_R were amplified from the human GPCR cDNA library (hORFeome database 8.1), and the third intracellular loop (ICL3) of each Ado receptor was swapped with the ICL3 of GRAB_NE_ (*2*). The swapping sites on A_2A_R and the amino acid composition between A_2A_R and ICL3 of GRAB_NE_ were then screened. The plasmids used to express the GRAB_Ado_ sensors in neurons or astrocytes were cloned into the pAAV vector using the human synapsin promoter (hSyn) or GfaABC1D promoter, respectively. The plasmids carrying subcellular compartment markers were cloned by fusing PSD95-RFP and synaptophysin-RFP into the pDest vector. To characterize signaling downstream of the GRAB_Ado_ sensors, constructs expressing GRAB_Ado_-SmBit and A_2A_R-SmBit were derived from β_2A_R-SmBit (*3*) using a BamHI site incorporated upstream of the GGSG linker. The LgBit-mGs plasmid (*3*) was a gift from Dr. Nevin A. Lambert. The plasmid encoding the cAMP sensor PinkFlamindo was generated using the published sequence (*4*). The plasmid encoding the glutamate sensor R^ncp^-iGluSnFR (*5*) was a gift from Dr. Robert Campbell, and the plasmid encoding TeNT was a gift from Dr. Peng Cao.

### Cell cultures

HEK293T cells and HeLa cells were cultured as described previously (*6*). In brief, HEK293T cells and HeLa cells were cultured at 37°C in 5% CO_2_ in DMEM (Biological Industries) supplemented with 10% (v/v) fetal bovine serum (FBS, Gibco) and 1% penicillin-streptomycin (Gibco).

Rat and mouse primary neurons were prepared from 0-day old (P0) pups (male and female, randomly selected). Cortical and hippocampal neurons were dissociated from the dissected brains in 0.25% Trypsin-EDTA (Gibco) and plated on 12-mm glass coverslips coated with poly-D-lysine (Sigma-Aldrich) in neurobasal medium (GIBCO) containing 2% B-27 supplement (Gibco), 1% GlutaMAX (Gibco), and 1% penicillin-streptomycin (Gibco). Based on glial cell density, after approximately 3 days in culture (DIV 3) cytosine β-D-arabinofuranoside (Sigma) was added to the hippocampal cultures in a 50% growth media exchange at a final concentration of 2 µM. Primary astrocytes were prepared as previously described (*7*). In brief, the hippocampi were dissected from P0 rat pups, and the cells were dissociated with trypsin digestion for 10 mins at 37°C and plated on a poly-D-lysine–coated T25 flask. The plating and culture media contained DMEM supplemented with 10% (v/v) FBS and 1% penicillin-streptomycin. The next day and every 2 days thereafter, the medium was changed. At DIV 7-8, the flask was shaken on an orbital shaker at 180 rpm for 30 min, and the supernatant containing the microglia was discarded; 10 ml of fresh astrocyte culture medium was then added to the flask, which was shaken at 240 rpm for ≥6 h to remove oligodendrocyte precursor cells. The remaining astrocytes were dissociated with trypsin and plated on 12-mm glass coverslips containing culture medium. Both the neurons and astrocytes were cultured at 37°C in 5% CO_2_.

### Mice

All animals were family- or pair-housed in a temperature-controlled room with a 12-h/12-h light/dark cycle. All protocols for animal surgery and maintenance were approved by the Animal Care and Use Committees at Peking University, the Chinese Academy of Sciences (CAS), and the National Institute of Biological Sciences, and were performed in accordance with the guidelines established by US National Institutes of Health. Both male and female mice (P0 pups for primary neuron cultures and adults >6 weeks for acute slices and *in vivo* experiments) were used for experiments. Nt5e (CD73) knockout mice were obtained from Jackson Laboratory (stock no. 018986).

### AAV virus preparation

The following AAV viruses were used to infect cultured neurons and/or astrocytes and for *in vivo* expression: AAV2/9-hSyn-Ado1.0, AAV2/9-hSyn-Ado1.0mut, AAV2/9-hSyn-R^ncp^-iGluSniFR, AAV2/9-GfaABC1D-Ado1.0, AAV2/9-hSyn-Ado1.0med, and AAV2/9-CAG-EBFP2-iP2A-TeNT (packaged at Vigene Biosciences); AAV9-CaMKIIα-hM3Dq-mCherry, AAV9-GfaABC1D-NES-jRGECO1a (packaged at BrainVTA Wuhan); and AAV2/9-hSyn-ChrimsonR-tdTomato (packaged at Shanghai Taitool Bioscience).

### Expression of GRAB_Ado_ in cultured cells and *in vivo*

HEK293T cells and HeLa cells were plated on 12-mm glass coverslips in 24-well plates and grown to 60-80% confluence for transfection. Cells were transfected using a mixture containing 1 μg DNA and 1 μg PEI for 4-6 h and imaged 24-48 h after transfection. For screening, cells expressing candidate GRAB_Ado_ sensors were plated in 96-well plates (PerkinElmer).

To achieve the sparse expression of Ado1.0 in neurons shown in Fig. 6A and fig. S2A, DIV 7-9 neurons were transfected using the calcium phosphate method as described previously (*6*). In brief, neurons were transfected with a mixture containing 125 mM CaCl_2_, 2x HBS (pH 7.04-7.12), and 2 μg DNA; 2 h after transfection, the neurons were washed with pre-warmed 1x HBS (pH 6.80) to remove the DNA-calcium phosphate precipitate. The neurons were imaged ≥48 h after transfection. For diffuse *in vitro* expression, the viruses were added to cultured neurons and astrocytes at DIV 7-9 and characterized ≥48 h after infection. For physiological analyses, DIV ≥13 neurons and astrocytes were used.

For *in vivo* expression, adult (>6 weeks of age) mice were anesthetized either with isoflurane (RWD Life Science) inhalation or Avetin (500 mg/kg, Sigma, i.p. injection), placed in a stereotaxic frame, and AAVs were injected using a microsyringe pump (Nanoliter 2000 Injector, WPI). In the chemical-induced seizure model, AAV expressing hSyn-Ado1.0 or hSyn-Ado1.0mut was injected into the right hippocampal CA1 region using the following coordinates: AP: −2.2 mm relative to Bregma, ML: −1.6 mm relative to Bregma, and DV: 1.2 mm below the dura. In the acute hypoxia model, AAV expressing hSyn-Ado1.0 or hSyn-Ado1.0mut were injected bilaterally into the mPFC using the following coordinates: AP: +1.78 mm relative to Bregma, ML: ±0.25 mm relative to Bregma, and DV: 2.4 mm below the dura); AAV was also injected bilaterally to the NAc using the following coordinates: AP: +0.98 mm relative to Bregma, ML: ±1.13 mm relative to Bregma, and DV: 4.65 mm below the dura). All experiments were performed at least one week after virus injection.

### Luciferase complementation assay

The luciferase complementation assay was performed as previously described (*2, 3*). In brief, Ado1.0-SmBit, A_2A_R-SmBit, or no construct was co-transfected with LgBit-mGs in HEK293T cells. Forty-eight hours after transfection, the cells were washed with phosphate-buffered saline (PBS), harvested by trituration, and transferred to opaque 96-well plates containing diluted Ado-containing solutions ranging from 0.1 nM to 100 mM Ado. Furimazine (NanoLuc Luciferase Assay, Promega) was added to each well immediately prior to performing the measurements with Nluc.

### Confocal imaging of GRAB_Ado_ in cultured cells

GRAB_Ado_-expressing HEK293T cells, HeLa cells, cultured neurons, and cultured astrocytes were screened using an Opera Phenix high-content imaging system (PerkinElmer) and imaged using a Ti-E A1 inverted confocal microscope (Nikon).

For screening GRAB_Ado_ sensors, a 60x/1.15 NA water-immersion objective was used. To image GFP and RFP fluorescence, we used a 488-nm and 561-nm laser, respectively, for excitation, and a 525/50-nm and 600/30-nm emission filter, respectively, to collect the signals. HEK293T cells expressing GRAB_Ado_ sensors were first bathed in Tyrode’s solution containing (in mM): 150 NaCl, 4 KCl, 2 MgCl_2_, 2 CaCl_2_, 10 HEPES, and 10 glucose (pH 7.3-7.4) and imaged before and after the addition of 100 μM Ado, and the change in GRAB_Ado_ fluorescence (ΔF/F_0_) was calculated using the ratio of GFP intensity to RFP intensity.

For confocal imaging, the microscope was equipped with either a 20x/0.75 NA objective or a 10x/0.5 NA objective, a 405-nm laser, a 488-nm laser, and a 561-nm laser. A 450/25-nm, a 525/50-nm, and a 595/50-nm emission filter was used to collect the BFP, GFP, and RFP signals, respectively. GRAB_Ado_-expressing HEK293T cells, neurons, and astrocytes were perfused with Tyrode’s solutions containing the drug of interest in the imaging chamber. During imaging, the cultured cells were bathed or perfused in a chamber with Tyrode’s solution. Solutions containing adenosine (Sigma), ADP (Sigma), ATP (Sigma), SCH-58261 (Abcam), HENECA (Tocris), inosine (Sigma), adenine (Sigma), CdCl_2_ (Sigma), L-glutamate (Sigma), bradykinin (Sangon Biotech Shanghai), thrombin (Sigma), POM1 (Santa Cruz), S-(4-nitrobenzyl)-6-thioinosine (NBTI, Santa Cruz), dipyridamole (Santa Cruz), ω-Conotoxin-GVIA (Tocris), ω-Agatoxin IVA (Cayman) nimodipine (Cayman), and (±)-felodipine (Cayman) were delivered via a custom-made perfusion system or via bath application. The chamber was cleaned thoroughly with Tyrode’s solution and 75% ethanol between experiments. The GFP signal (e.g., the GRAB_Ado_ sensors and A_2A_R-EGFP) was collected using a 525/50-nm emission filter, the RFP signal (mCherry-CAAX, PinkFlamindo, Calbryte 590, and R^ncp^-iGlu) was collected using a 595/50-nm emission filter, and the BFP signal was recorded using a 450/25-nm emission filter. To measure kinetics, 100 μM Ado and 200 μM SCH-58261 were applied with rapid perfusion. Where applicable, cells were pre-loaded with the Ca^2+^ dye Calbryte 590-AM (AAT Bio) by incubation at 37°C for 40 min before imaging. High K^+^–containing Tyrode’s solution contained (in mM): 79 NaCl, 75 KCl, 2 MgCl_2_, 2 CaCl_2_, 10 HEPES, and 10 glucose (pH 7.3-7.4). For field stimulation, neurons were stimulated using platinum electrodes as previously described (*8*); except where indicated otherwise, the stimulation voltage was set at 80 V, and the duration of each stimulation pulse was typically set at 1 ms. All experiments were performed at room temperature.

### Preparation and fluorescence imaging of acute brain slices

Wild-type adult (42-56 days of age) C57BL/6N mice were anesthetized with an intraperitoneal injection of Avertin (500 mg/kg body weight) and placed in a stereotaxic frame for AAV injection using a microsyringe pump (Nanoliter 2000 Injector, WPI). For the data in fig. S8A-D, AAV expressing hSyn-Ado1.0med was injected into the hippocampal CA3 region using the following coordinates: AP: +1.8 mm relative to Bregma, ML: ±2.2 mm relative to Bregma, and depth: 1.5 mm below the dura. For the data in Fig. 6B-F and fig. S8E-H, AAVs expressing hSyn-Ado1.0med and hSyn-ChrimsonR-tdTomato (or CaMKIIα-hM3Dq-mCherry) were injected into the hippocampal CA1 region using the following coordinates: AP: +1.8 mm relative to Bregma, ML: ±1.0 mm relative to Bregma, and depth: 1.2 mm from the dura. For the data in fig. S7, AAVs expressing hSyn-Ado1.0med were injected into the mPFC region using the following coordinates: AP: +1.78 mm relative to Bregma, ML: ±0.25 mm relative to Bregma, and depth: 2.4 mm below the dura

Two to four weeks after virus injection, the mice were anesthetized with an intraperitoneal injection of Avertin (500 mg/kg body weight) and perfused with ice-cold oxygenated slicing buffer containing (in mM): 110 choline chloride, 2.5 KCl, 0.5 CaCl_2_, 7 MgCl_2_, 1.3 NaH_2_PO_4_, 25 NaHCO_3_, 10 glucose, 1.3 Na-ascorbate, and 0.6 Na-pyruvate. The brains were immediately removed and placed in ice-cold oxygenated slicing buffer. The brains were then sectioned into 200-μm thick slices using a VT1200 vibratome (Leica), and the slices were incubated at 34°C for at least 40 min in oxygenated artificial cerebrospinal fluid (ACSF) containing (in mM): 125 NaCl, 2.5 KCl, 2 CaCl_2_, 1.3 MgCl_2_, 1.3 NaH_2_PO_4_, 1.3 Na-ascorbate, 0.6 Na-pyruvate, 10 glucose, and 25 NaHCO_3_. For the data in fig. S8, the slices were transferred to an imaging chamber and placed under an FV1000MPE 2-photon microscope (Olympus) equipped with a 25x/1.05 NA water-immersion objective and a mode-locked Mai Tai Ti:Sapphire laser (Spectra-Physics). A 920-nm laser was used to excite Ado1.0med, and fluorescence was collected using a 495-540-nm filter. For electrical stimulation, a bipolar electrode (cat. number WE30031.0A3, MicroProbes for Life Science) was positioned near the CA1 stratum radiatum using fluorescence guidance. Fluorescence imaging and electrical stimulation were synchronized using an Arduino board with custom-written programs. All images collected during electrical stimulation were recorded at a frame rate of 2.8 frames/s, with 256×192 pixels per frame. The stimulation voltage was 4–8 V, and the duration of each stimulus was 1 ms. Drugs were applied to the imaging chamber by perfusion at a flow rate at 4 ml/min. For the data in Fig. 6B-F and fig. S7, Ado1.0med fluorescent signals were imaged with a 20× water immersion objective on an Olympus FV1000 microscope at the rate of 0.795 sec/frame with 320×320 pixels in each frame. For photo-stimulation, an optical fiber (200 μm core diameter, NA = 0.22) coupled to a diode-pumped solid-state 633 nm laser was submerged in ACSF and placed ∼500 μm from the imaging region.

### Fiber photometry and local field potential recordings in living mice

In the chemical-induced seizure model, 1 week after virus injection a cannula was implanted into the left hippocampal CA1 region using the following coordinates: AP: −2.2 mm relative to Bregma, ML: −1.6 mm relative to Bregma, and DV: 1.2 mm below the dura) and used for drug delivery; an optical fiber (Thorlabs, FT200UMT) and an electrode were inserted to the right hippocampal CA1 region using same coordinates for virus injection and was used to record the optical signals and local field potentials (LFPs). At least 1 week after surgery, kainic acid (KA, Cayman, ∼0.25 μg in 0.1 μl saline) was injected through the cannula to induced acute seizures. To record GRABAdo fluorescence, a 459-Hz sinusoidal blue LED light (CLED_465, ∼30 μW, Thorlabs) was bandpass-filtered (passing band: 460-490 nm) and delivered to the brain to excite the sensor; the emitted light then traveled back through the same optical fiber, was bandpass-filtered (passing band: 500-550 nm), detected using a Femtowatt Silicon Photoreceiver (Newport, 2151), and recorded using a real-time processor (RZ2, TDT). The envelope of the 459-Hz signal that reflects the intensity of the fluorescence signals was extracted in real-time using a custom TDT program.

In the acute hypoxia model, following virus injection an optical fiber (OD: 230 μm, NA: 0.37; Shanghai Fiblaser) was placed in a ceramic ferrule and advanced toward the mPFC and NAc through the craniotomy. The ceramic ferrule was affixed with a skull-penetrating M1 screw and dental acrylic. The mice were then housed individually for at least 1 week to recover. To induce acute hypoxia, the mouse was placed in a plastic cylindrical chamber (diameter: 15 cm, height: 15 cm) and 99.99% N_2_ was infused through the bottom of the chamber at a flow rate of 4.5 L/min for 50 s. To record GRAB_Ado_ fluorescence, a 470-nm excitation light was reflected by a dichroic mirror (#67079, Edmund) and focused using a 20x/0.4 objective. A 2-meter long custom patch cord containing four bundled optical fibers (OD: 230 mm, NA: 0.37) was used to deliver the excitation light and collect the emission light. To minimize photobleaching at the tip of the optical fiber, the laser power was adjusted to a low level (0.01– 0.02 mW). The GRAB_Ado_ fluorescence signal was bandpass-filtered (MF525-39, Thorlabs) and captured as a series of images using a CMOS camera (DCC3240M, Thorlabs). The fluorescence intensity profile of each channel was then extracted by averaging the pixel values in the relevant ROIs. The acquisition frequency was 50 Hz, and the data were acquired and stored on a computer using a custom-written LabView program.

### Histology

To verify the expression of viruses and proper placement of the optical fibers, the mice were deeply anesthetized and immediately perfused with 0.1 M PBS, followed by 4% paraformaldehyde (PFA). The brains were removed and post-fixed overnight in 4% PFA before dehydration using a 30% (w/v) sucrose solution. The brains were embedded in OCT compound (Thermo Scientific) and cut into 50-μm sections using a cryostat (Thermo Scientific). The brain sections were stored at −20°C until further processing. For histology examination, the brain sections were washed in PBS and coverslipped with VECTASHIELD mounting media containing DAPI (Vector Laboratories). The fluorescence images were captured using an epifluorescence microscope (VS120, Olympus).

### Data analysis

Imaging data from cultured cells were first processed using ImageJ software (NIH); traces were generated using OriginPro 2019, and pseudocolor images were generated using ImageJ.

To analyze the photometry data in the chemical-induced seizure model, we first binned the raw data into 1-Hz bins (i.e., down-sampled by 1000) and then subtracted background autofluorescence. We calculated ΔF/F_0_ using a baseline obtained by fitting the autofluorescence-subtracted data with a 2^nd^-order exponential function. In the acute hypoxia model, we calculated the change in fluorescence essentially as described previously (*9*); in brief, we derived ΔF/F_0_ by calculating [(F-F_0_)/F_0_], where F_0_ is the average fluorescence signal intensity over a 2-s window prior to N_2_ infusion.

Except where indicated otherwise, all summary data are presented as the mean ± SEM. Group differences were analyzed using the paired or unpaired Student’s *t*-test or one-way repeated measures ANOVA; in all figures, \**p*<0.05; \*\**p*<0.01, \*\*\**p*<0.001, and n.s., not significant (*p*>0.05).

## References

1. A. Drury, A. v. Szent-Györgyi, The physiological activity of adenine compounds with especial reference to their action upon the mammalian heart 1. The Journal of Physiology 68, 213–237 (1929).

2. J.-F. Chen, H. K. Eltzschig, B. B. Fredholm, Adenosine receptors as drug targets—what are the challenges? Nat Rev Drug Discov 12, 265 (2013).

3. R. B. Dias, D. M. Rombo, J. A. Ribeiro, J. M. Henley, A. M. Sebastiao, Adenosine: setting the stage for plasticity. Trends Neurosci 36, 248–257 (2013).

4. G. Hasko, J. Linden, B. Cronstein, P. Pacher, Adenosine receptors: therapeutic aspects for inflammatory and immune diseases. Nat Rev Drug Discov 7, 759–770 (2008).

5. R. A. Cunha, How does adenosine control neuronal dysfunction and neurodegeneration. Journal of Neurochemistry 139, 1019–1055 (2016).

6. J. Feng et al., A Genetically Encoded Fluorescent Sensor for Rapid and Specific In Vivo Detection of Norepinephrine. Neuron 102, 745–761 e748 (2019).

7. T. Patriarchi et al., Ultrafast neuronal imaging of dopamine dynamics with designed genetically encoded sensors. Science 360, eaat4422 (2018).

8. F. Sun et al., A Genetically Encoded Fluorescent Sensor Enables Rapid and Specific Detection of Dopamine in Flies, Fish, and Mice. Cell 174, 481–496 e419 (2018).

9. M. Jing et al., A genetically encoded fluorescent acetylcholine indicator for in vitro and in vivo studies. Nat Biotechnol 36, 726–737 (2018).

10. Y. Lee, A. Messing, M. Su, M. Brenner, GFAP promoter elements required for region-specific and astrocyte-specific expression. Glia 56, 481–493 (2008).

11. Y. H. Raol, A. R. Brooks-Kayal, in Progress in molecular biology and translational science. (Elsevier, 2012), vol. 105, pp. 57–82.

12. L. Weltha, J. Reemmer, D. Boison, The role of adenosine in epilepsy. Brain research bulletin 151, 46–54 (2019).

13. H. Winn, R. Rubio, R. Berne, Brain adenosine concentration during hypoxia in rats. American Journal of Physiology-Heart Circulatory Physiology 241, H235–H242 (1981).

14. W. Peng et al., Adenosine dynamics during sleep-wake cycles. Submitted, (2020).

15. D. Lovatt et al., Neuronal adenosine release, and not astrocytic ATP release, mediates feedback inhibition of excitatory activity. Proc Natl Acad Sci U S A 109, 6265–6270 (2012).

16. E. D. Martín et al., Adenosine released by astrocytes contributes to hypoxia-induced modulation of synaptic transmission. Glia 55, 36–45 (2007).

17. G. Burnstock, Purinergic signalling and disorders of the central nervous system. Nat Rev Drug Discov 7, 575 (2008).

18. M. J. Wall, N. Dale, Neuronal transporter and astrocytic ATP exocytosis underlie activity-dependent adenosine release in the hippocampus. J Physiol 591, 3853–3871 (2013).

19. F. Corti et al., Adenosine is present in rat brain synaptic vesicles. Neuroreport 24, 982–987 (2013).

20. Z. Zhang, K. T. Nguyen, E. F. Barrett, G. David, Vesicular ATPase inserted into the plasma membrane of motor terminals by exocytosis alkalinizes cytosolic pH and facilitates endocytosis. Neuron 68, 1097–1108 (2010).

21. J. Wu et al., Genetically encoded glutamate indicators with altered color and topology. ACS Chem Biol 13, 1832–1837 (2018).

22. G. G. Schiavo et al., Tetanus and botulinum-B neurotoxins block neurotransmitter release by proteolytic cleavage of synaptobrevin. Nature 359, 832 (1992).

23. M. Patterson, E. Szatmari, R. Yasuda, AMPA receptors are exocytosed in stimulated spines and adjacent dendrites in a Ras-ERK–dependent manner during long-term potentiation. Proc Natl Acad Sci U S A 107, 15951–15956 (2010).

24. H. Vacher, D. P. Mohapatra, J. S. Trimmer, Localization and targeting of voltage-dependent ion channels in mammalian central neurons. Physiological Reviews 88, 1407–1447 (2008).

25. J. W. Hell et al., Identification and differential subcellular localization of the neuronal class C and class D L-type calcium channel alpha 1 subunits. Journal of Cell Biology 123, 949–962 (1993).

26. D. B. Wheeler, A. D. Randall, R. W. Tsien, Roles of N-type and Q-type Ca2+ channels in supporting hippocampal synaptic transmission. Science 264, 107–111 (1994).

27. F. A. Dodge, R. Rahamimoff, Co-operative action of calcium ions in transmitter release at the neuromuscular junction. The Journal of Physiology 193, 419–432 (1967).

28. R. Fernandezchacon et al., Synaptotagmin I functions as a calcium regulator of release probability. Nature 410, 41–49 (2001).

29. B. B. Fredholm, Adenosine, an endogenous distress signal, modulates tissue damage and repair. Cell Death & Differentiation 14, 1315–1323 (2007).

30. M. M. Halassa et al., Astrocytic Modulation of Sleep Homeostasis and Cognitive Consequences of Sleep Loss. Neuron 61, 213–219 (2009).

31. J. M. Brundege, T. V. Dunwiddie, Modulation of Excitatory Synaptic Transmission by Adenosine Released from Single Hippocampal Pyramidal Neurons. Journal of Neuroscience 16, 5603–5612 (1996).

32. O. J. Manzoni, T. Manabe, R. A. Nicoll, Release of adenosine by activation of NMDA receptors in the hippocampus. Science 265, 2098–2101 (1994).

33. D. Brambilla, D. Chapman, R. W. Greene, Adenosine Mediation of Presynaptic Feedback Inhibition of Glutamate Release. Neuron 46, 275–283 (2005).

34. Z. Wu, Y. Li, New frontiers in probing the dynamics of purinergic transmitters in vivo. Neuroscience Research 152, 35–43 (2020).

## References

1. D. G. Gibson et al., Enzymatic assembly of DNA molecules up to several hundred kilobases. Nature Methods 6, 343–345 (2009).

2. J. Feng et al., A Genetically Encoded Fluorescent Sensor for Rapid and Specific In Vivo Detection of Norepinephrine. Neuron 102, 745–761 e748 (2019).

3. Q. Wan et al., Mini G protein probes for active G protein–coupled receptors (GPCRs) in live cells. Journal of Biological Chemistry 293, 7466–7473 (2018).

4. K. Harada et al., Red fluorescent protein-based cAMP indicator applicable to optogenetics and in vivo imaging. Scientific reports 7, 1–9 (2017).

5. J. Wu et al., Genetically encoded glutamate indicators with altered color and topology. ACS Chem Biol 13, 1832–1837 (2018).

6. F. Sun et al., A Genetically Encoded Fluorescent Sensor Enables Rapid and Specific Detection of Dopamine in Flies, Fish, and Mice. Cell 174, 481–496 e419 (2018).

7. S. Schildge, C. Bohrer, K. Beck, C. Schachtrup, Isolation and culture of mouse cortical astrocytes. JoVE, e50079 (2013).

8. Y. Li, R. W. Tsien, pHTomato, a red, genetically encoded indicator that enables multiplex interrogation of synaptic activity. Nature Neuroscience 15, 1047 (2012).

9. Y. Li et al., Hypothalamic circuits for predation and evasion. Neuron 97, 911–924. e915 (2018).

